# I TRIED A BUNCH OF THINGS: THE DANGERS OF UNEXPECTED OVERFITTING IN CLASSIFICATION

**DOI:** 10.1101/078816

**Authors:** Michael Powell, Mahan Hosseini, John Collins, Chloe Callahan-Flintoft, William Jones, Howard Bowman, Brad Wyble

**Affiliations:** Manada Technology LLC; Computing Department, University of Kent; Physics Department, Penn State University; Army Research Lab, Aberdeen Proving Grounds; School of Psychology, University of Birmingham; Psychology Department, Penn State University

## Abstract

Machine learning is a powerful set of techniques that has enhanced the abilities of neuroscientists to interpret information collected through EEG, fMRI, and MEG data. With these powerful techniques comes the danger of *overfitting of hyper-parameters* which can render results invalid, and cause a failure to generalize beyond the data set. We refer to this problem as *‘over-hyping’* and show that it is pernicious despite commonly used precautions. In particular, over-hyping occurs when an analysis is run repeatedly with slightly different analysis parameters and one set of results is selected based on the analysis. When this is done, the resulting method is unlikely to generalize to a new dataset, rendering it a partially, or perhaps even completely spurious result that will not be valid outside of the data used in the original analysis. While it is commonly assumed that cross-validation is an effective protection against such spurious results generated through overfitting or overhyping, this is not actually true. In this article, we show that both one-shot and iterative optimization of an analysis are prone to over-hyping, despite the use of cross-validation. We demonstrate that non-generalizable results can be obtained even on non-informative (i.e. random) data by modifying hyper-parameters in seemingly innocuous ways. We recommend a number of techniques for limiting over-hyping, such as lock-boxes, blind analyses, pre-registrations, and nested cross-validation. These techniques, are common in other fields that use machine learning, including computer science and physics. Adopting similar safeguards is critical for ensuring the robustness of machine-learning techniques in the neurosciences.

## INTRODUCTION

Computers have revolutionized approaches to data analysis in psychology and neuroscience, effectively allowing one to interpret not only the neural correlates of cognitive processes, but also the information content that is represented in the brain through the use of machine learning. However, with these new and powerful tools come new dangers. Machine learning algorithms allow a pattern classifier to weave many subtle threads of information together to detect hidden patterns in a massive data set, e.g. to determine from MEG data whether someone is currently viewing a building or an animal (Cichy, Pantavis & Oliva 2014). However, these pattern classifiers are essentially black boxes to their human operators, as they create complex mappings between features and outputs that exceed one’s ability to comprehend. Consequently, when using such algorithms, one can unintentionally overfit to the data, even when performing analyses that intuitively seem correct. This article seeks to provide a clear understanding of the dangers of a particular form of overfitting we refer to as over-hyping through example, and describes tools to limit and detect it. We feel that a thorough understanding of the error introduced through over-hyping is crucial, since this error is easy to commit yet difficult to detect. Furthermore, while there has been a lot of discussion of problems of circularity and inflated effects in neuroscience analyses (e.g. Kriegeskorte, Simmons, Bellgowan & Baker 2009; Vul, Harris, Winkielman & Pashler 2009; Eklund, Nichols, Anderson & Knutsson 2015; Brooks, Zoumpoulaki & Bowman, 2017), machine learning algorithms are so effective that they provide dangers above and beyond those that have been discussed. Optimization of hyper-parameters is a common practice in the machine learning literature (Bouthillier & Varoquaux 2020) and it is difficult to determine how the data were treated during the optimization process. Importantly, as will be demonstrated below, *the technique of cross-validation, often employed as a safeguard against overfitting, is not entirely effective at ensuring generalizability*. We suspect that the incidence of accidental overfitting errors in the literature could be substantial already, and may increase as machine learning methods increase in popularity.

Overfitting is the optimization of a model on a given data set such that performance improves on the data being evaluated but remains constant or degrades on other similar data. This ‘other’ data can be referred to as *out-of-sample*, meaning that it is outside of the data that was used to train and evaluate the classifier. In other words, if one were to develop a machine learning approach on one data set and then apply the same algorithm to a second set of data drawn from the same distribution, performance might be much worse than on the original set of data. This is a severe problem, because models that cannot generalize to out-of-sample data have little to say about brain function in general: Their results are valid only on the data set used to configure the classifier, are tuned to the specific pattern of noise in the data, and are unlikely to be replicated on any other set. One of the earlier and more startling examples of overfitting was performed by Freedman (1983), where he showed—with high statistical significance—that a regression model could be used to find a strong relationship between independent random variables drawn from a standard normal distribution (which have no real relationship whatsoever).

Problems with overfitting are by no means new to science: High-energy physics has had a number of high-profile false discoveries, some of which were the result of overfitting an analysis to a particular data set. For accounts of some of these in the light of current experimental practice, see Harrison (2002) and Dorigo (2015). The similarities between data analysis in high-energy physics and modern neuroscience are striking: both fields have enormous quantities of data that need to be reduced to discover signals of interest. As such, it is useful and common to apply cuts to the data, i.e. to restrict analysis to certain regions of interest (ROI), as is common to the analysis of fMRI and EEG data. Because the purpose of the cuts is to enhance a signal of interest, there is a danger that the choice of a cut made on the basis of the data being analyzed (and on the basis of the desired result) may create apparent signal where none actually exists. Furthermore, when making measurements in high-energy physics and neuroscience, complicated apparatuses are often used, and analyses typically contain an extremely sophisticated set of software algorithms. Optimization, and debugging of the analysis chain has further potential for creating an apparent measurement of a non-existent effect. Making these cuts and other analysis optimizations require many decisions that are often necessary to process the data, and yet present grave dangers to the generalizability of the results.

To better understand the principles of this conundrum, we rely on a commonly used distinction between parameters and hyper-parameters. In the context of machine learning, we use the term parameter to refer to aspects of the analysis that are directly driven by the data. For example when training a support-vector-machine, the classification function learnt would be defined by a set of parameters. They are learnt through direct contact with the data. Hyper-parameters, on the other hand, refer to values that are configured to improve model performance. These will include, but are not necessarily limited to the following: artifact rejection criteria, feature selection (i.e. electrodes or ROIs in the brain), filter settings, control parameters of classifiers (e.g. choice of kernels, setting of regularisation parameters), and even choice of classifier (e.g. SVM vs. LDA vs. random forests vs naïve Bayes). These are settings and choices that could, at least in principle, apply across a class of data sets.

In this context, we propose the term *over-hyping* (i.e. overfitting of hyperparameters) as a specific case of (typically unintentional) overfitting through adjustment of analysis hyper-parameters in conjunction with consultation of analysis results. We suggest that over-hyping is a fairly widespread and poorly understood problem in the neurosciences, particularly because the field utilizes relatively expensive and time consuming data collection practices (unlike the field of machine-vision, for example, which has a plentiful source of new data). Over-hyping is likely to be a problem in any field with similar cost constraints on data collection. Indeed, because of several high-profile false discoveries, high-energy physics has already gone through a replicability crisis, and has had to rearrange its methods to deal with the consequences. A classic case, which was a big wake-up call, was the so-called split-A2 from Chikovani et al. (1967). Had this effect been genuine, it would have engendered a theoretical revolution, but when more data became available, the effect disappeared; see Harrison (2002) for a recent view. It appeared that inappropriate selection of data was the culprit, which is a form of over-hyping.

To prevent such cases, the high-energy physics community has adopted several conventions and methods in the analysis and interpretation of data. Of these changes, the one that is most relevant to over-hyping is called *blind analysis*, referring to a technique in which analysis optimization occurs without consulting the dependent variable of interest (e.g. Klein & Roodman 2005). Since the optimization algorithm is blind to the result of interest, over-hyping is at least difficult, if not impossible. At the end of this paper, we will discuss several preventative solutions, including blind analysis. Unlike physics, while related issues have been discussed in the literature (Kriegeskorte et al 2009; Brooks et al 2017), the neuroscience field has not yet fully responded to the dangers of over-hyping when complex analyses are used, which increases the potential of false findings and presents a major barrier to the replicability of the literature.

### CROSS-VALIDATION DOES NOT PREVENT OVER-HYPING WHEN RE-USED ON THE SAME DATA SET

In the neuroscience literature, a method that is typically employed to prevent over-fitting is cross-validation, in which data are repeatedly partitioned into two non-overlapping subsets. In each iteration, classifiers are trained on one set and tested on the other and the results of multiple iterations are averaged together. Because the training and testing sets are disjoint in each iteration, the average performance on the test sets is believed to provide an unbiased estimate of classifier performance on out-of-sample data. However, this is only true as long as one important restriction is obeyed: After performing cross-validation, the analysis chain used to classify the data must *not* be modified to obtain higher performance *on that same data*. Reusing the same data to optimize analysis parameters can induce over-hyping, *even if cross-validation is used at each iteration*.

The reason that over-hyping can occur despite cross-validation is that all data sets are composed of a combination of signal and noise. The signal is the portion of the data containing the useful information that one would like the machine learning classifier to discover, while the noise includes other sources of variability. However, when an analysis is optimized on a given data set after viewing the results, the choice of hyper-parameters can be influenced by how the noise affected the classification accuracy. Consequently, while the optimization improves classification accuracy on this data set, performance may remain constant or even worsen on a completely distinct set of data, because (in a statistical sense) its noise is not shared with the data driving the optimization (Figure 1). In other words, the analysis would not replicate on a distinct dataset even if the sampling conditions were identical.

**Figure 1.**
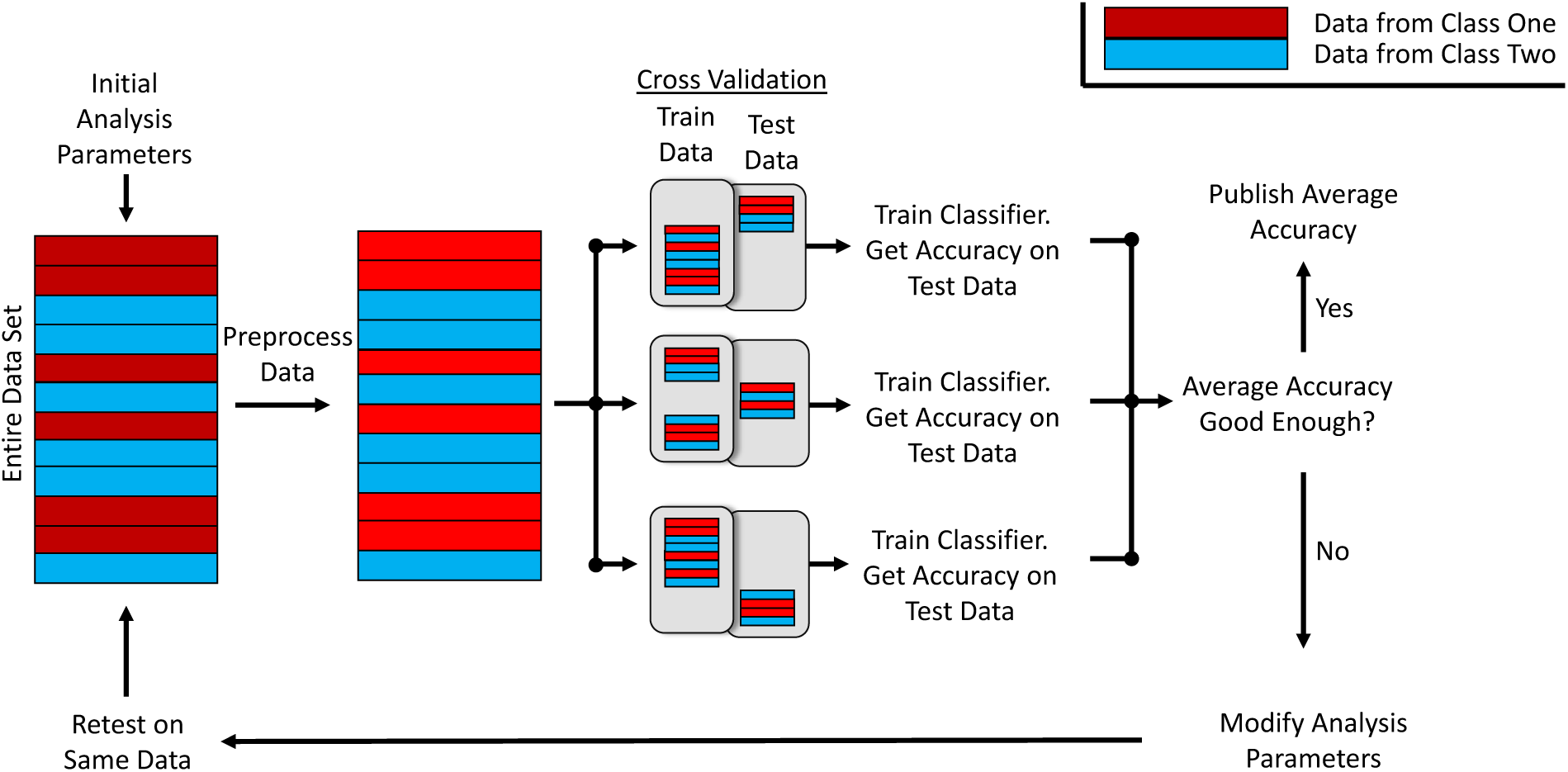
An example of how over-hyping can be induced by modifying hyper-parameters after evaluating a system through cross-validation. The feedback loop allowing hyper-parameters to be adjusted after viewing the results provides a route for analysis decisions to be made in response to the noise in the data set, despite the separation of data into training/testing sets.

The possibility that cross-validation does not prevent over-hyping, is well known in the machine learning and machine vision communities (Domingos 2012), which are taking increasing care to avoid the problem. For example, machine-learning competitions on websites such as Kaggle.com provide contestants with sample data on which to optimize their models. The final evaluation of the contestants is performed on a different data set that is either held as confidential by the sponsoring organization, or is released only a few days before the end of the competition. Contestants who access the data more often than the rules permit are disqualified, their organizations can be barred from future competitions and in one recent high-profile case a lead scientist was fired (Markoff 2015).

In writing this paper, we share the experience of our colleagues in the physics and computer science disciplines so as to encourage more rigorous standards of machine learning before a replicability crisis in neuroscience machine learning unfolds. It is not our intent to call out specific examples of bad practice in the literature, although in our informal survey of the neuroscience classification literature we found very few cases in which the appropriate precautions had been documented (i.e. some variety of blind, nested cross-validation or lock-box analysis, which will be described below). Without these precautions it is impossible to determine from a paper’s methods whether overfitting of hyper-parameters occurred. This is concerning because it is unlike the fallacy of double-dipping (Kriegeskorte, Simmons, Belgowan & Baker 2009), which is clearly discernible in the methods section of the paper. The difficulty in identifying cases of over-hyping is that it would have occurred during the optimization of the analysis, and these steps are typically omitted from the methods. Another issue that we observe in the literature is inconsistent terminology, which makes it harder to understand exactly what was done (e.g. Ng (1997) and Varoquaux (2017) use different definitions of ‘test set’). To help clarify terminology, we offer a table describing common terms and descriptions of what they are typically taken to mean (Table 1). We suggest a new term, the *Lock box*, which refers to a set of data that is held-out from the optimization process for verification and should not be consulted until the method’s hyper-parameters have been completely determined. The term *held-out* data set is sometimes taken to mean this, but that term is also used inconsistently and is easy to misinterpret. The term *lock-box* more clearly indicates the importance of holding the data in an inaccessible reserve. More will be said about this below. Next, we provide clear examples of over-hyping despite use of cross-validation using a sample of EEG data recorded from our own lab. We use real data instead of simulated data, to ensure that the noise reflects the genuine variability typically found in similar datasets. In the experiment, EEG data were collected while subjects tried to detect visually presented targets.

**Table 1.** The terminology used in this and other papers are defined in this table.

| Term | Definition |
| --- | --- |
| Machine Learning | Machine learning is the use of semi-automated fitting algorithms to discern patterns in data. Typically, machine learning algorithms are trained with labeled data from two or more classes, and are then used to predict which class a new and unlabeled data item belongs to. Examples of machine learning algorithms are support vector machine (SVM) classifiers, random forest models, and naive Bayes classifiers. |
| Cross-Validation | A technique commonly used to evaluate classification performance that repeatedly divides the data into two subsets (each division is a **fold**), one of which is used to train a classifier, the other being used to test it. |
|  | Performance is taken as the average across all folds. See the supplemental for a more thorough description. |
| Training and Testing sets | These terms generally refer to the two subsets of data used during cross-validation. However, the term test-set is sometimes used to refer to data that has been set-aside for later evaluation. We advise against that usage for the sake of consistency. |
| Nested Cross-Validation | Nested cross-validation is a generalization of cross-validation in which the data are now partitioned into N outer sets/ folds. Each of these folds provides an outer hold-out set, and an inner set, on one outer cycle. On each such outer cycle, cross-validation is performed on the inner set. The benefit of the nested approach is that it provides a reliable assessment of overfitting of hyper-parameters. See the supplemental for a more thorough description. |
| Lock-box | We introduce the term lock-box to mean a subset of data that are removed from the analysis pipeline at the very start of optimization and not accessed until all hyper-parameter adjustments and training have been completed. |
| Hyper-parameter | Hyper-parameters are a kind of parameter whose values are adjusted either by hand or by algorithms to improve model performance (e.g. weights of electrodes, regions of interest, SVM model kernel functions, classification model types). They are distinct from other parameters, whose values are set during classifier training (e.g. the linear function that results from training a least squares model or the classification function that results from training a Support Vector Machine classifier). |

The first example shown here is a one-shot hyper-parameter adjustment, in which 40 variations of machine classification are tested using cross-validation on a set of randomly scrambled data (i.e. there is no signal). By taking the most favorable result from these 40 variations, we evaluate whether a spurious result can be obtained by making a single hyperparameter choice despite cross-validation. The one-shot analysis shows a significant classification effect using conventional temporal-generalization analyses that are currently favored by the neuroimaging classification community (e.g. King & Dehaene 2014 TINS, King.. Dehaene PLOSONE 2014, Cichy et al., 2014). The illusory effect obtained by over-hyping, though small, would provide erroneous evidence of target discrimination in the EEG data over long periods of time, which is commonly taken as evidence that a neural correlate of working memory has been measured. A comparison to a Lock Box data set (i.e. data that were not consulted during analysis optimization) reveals that there is no reliable classification of target presence by this chosen set of hyperparameters, which, due to analyzed data sets not containing signal, reflects the expected outcome.

In the second example, we show a more extreme case of overhyping. Hyper-parameters were iteratively optimized to eliminate some features of the data set through a genetic algorithm using cross-validation at each step. Performance was compared to lock-box data that were set aside and not used in the genetic algorithm’s fitness function. Performance was shown to improve on the data on which the classifiers were optimized, but not on the lock-box data. Note that highly robust over-hyping was relatively easy to evoke, despite the use of cross-validation. The obtained results, presented below, demonstrate that classifiers can easily be over-hyped to obtain performance that will not generalize to set-aside or out-of-sample data.

## METHODS

### EEG METHODS

The simulations presented below were performed on EEG data, which was collected from rapid serial visual presentation (RSVP; Experiment 3 of Callahan-Flintoft, Chen and Wyble 2018; see supplemental for comprehensive methods). Subjects viewed a series of changing letters, presented bilaterally, updating at intervals of 150ms, and were tasked with reporting the one or two digits that would appear on each trial. For this analysis, we selected the trials containing either a single digit, or two digits presented in sequence separated by 600msand attempted to classify for each trial, whether one or two digits had been presented. However, the trial labels were randomly shuffled within subjects to obscure any actual effect of this manipulation. During each trial, EEG was recorded at 32 electrode sites and according to the standard 10-20 system. It was further bandpass filtered from 0.05 – 100 Hz, originally sampled at 500 Hz and down-sampled offline to 125 Hz for the present analysis. For further details on pre-processing and artifact rejection, see the EEG recordings section of experiment one in the original paper (Callahan-Flintoft et al., 2018). The original study excluded one subject due to an insufficient number of trials after artifact rejection. We decided to exclude an additional subject, choosing the one with the least number of trials, in order to be able to split the data into two equal parts for parameter optimization and lock box, detailed below. The final number of subjects was 24. Finally, the original study divided the data based on the visual hemifield in which the target stimulus was presented and only included trials in which correct responses were provided. We collapsed the data across hemifields and included all trials regardless of accuracy to increase the number of available trials per subject.

### OVERHYPING DUE TO KERNEL SELECTION DESPITE CROSS-VALIDATION

The main analysis used was a temporal generalisation, which determines whether a classifier trained at one point in time relative to stimulus onset is able to classify at other time points. Such analyses have been used to examine whether memory representations are stable over time in working memory research (e.g. Dehaene & King 2014).

A total of 1038 iterations was performed. In each one the trials were randomly shuffled within each subject. This needs to be stressed: all analyses were exclusively performed on null-data. Hence, any systematic improvements above chance performance must be due to over-hyping. Also, the data were randomly split into two equal parts of 12 subjects. One set was for parameter optimization (PO) and the other was the lock box (LB) set. The data were reshuffled into new PO and LB sets approximately every 15 iterations to ensure that any effects were not subject-specific. Our temporal generalisation analyses used functions of the MVPA-Light toolbox (M. Treder).

For each of the 1038 iterations randomized PO & LB data sets were created. 40 configurations of hyperparameters were used in each iteration to generate temporal generalisation maps to determine which configuration had produced the most desired outcome classifying the random PO data set. The 40 configurations were derived from 4 different classifiers: support vector machines (SVM) with three different kernels (linear, polynomial (order of 2); radial basis function (RBF)) and a linear discriminant analysis (LDA). Additionally, the extent of regularization was varied through 10 choices for each classifier. For SVMs, the C parameter took values of 0.0001, 0.0007, 0.0059, 0.0464, 0.03593, 2.7825, 21.5443, 166.81, 1291.5496 and 10000. The choice of C values was inspired by (and equal to) the search space of MVPA-Light’s default regularization search for SVMs. For LDA, candidate lambdas were 1, 0.88, 0.77, 0.66, 0.55, 0.44, 0.33, 0.22, 0.11 and 0. Temporal generalisation analyses were performed using 5-fold cross-validation.

These 40 candidate configurations competed in each of the 1038 iterations of the analysis, which we call *PO Competition* as they represent a competition between Parameter Optimizations. This PO competition was decided using a measure we call classification mass (*C-Mass*), which was computed on group-average temporal generalization maps (i.e. averages of 5-fold cross-validated single-subject maps). C-Mass reflects the average deflection from chance-level classification across the entire temporal generalization map. It was computed by first subtracting chance-level classification (Area Under the Curve/AUC of 0.5) from all AUC values of a given map, then squaring these values and finally taking the average of the entire map^1^.

For each of the 1038 iterations, hyperparameter configurations that led to maximum C-Mass classifying the PO data set were selected as the respective winner of the PO competition. The winners of this competition were then used to assess the degree of over-hyping by comparing them to the LB set.

We selected one of the 1038 iterations to demonstrate how manual selection could produce *what appears to be* a theoretically meaningful result in a temporal generalization map (Figure 2). Evaluating the efficacy of different hyperparameter configurations on the same dataset can be considered an analog of an exploratory analysis in which an analyst runs a series of cross-validated pilot analyses and stops on finding one that is theoretically suitable. In the working memory literature, it is considered important that a classifier is able to decode the condition label after the stimulus has disappeared. In our manually selected case, the winning PO map can be regarded theoretically suitable as it appears to exhibit this property whereby the accuracy remains well above chance for a substantial period of time after the target onset (see the supplemental for a randomly selected set of 9 additional iterations). However, this effect is demonstrably spurious, since the trial labels were all randomly shuffled. To demonstrate that this observed pattern is due to overhyping and not a general property of our analysis, we also present the results of the same analysis configuration for the companion LB set as well as of an alternative configuration for the same PO set. It is clear that the pattern observed in the winning PO map does not generalize either to another data set drawn from the same population using the same kernel configuration (the LB set) or to a reanalysis of the same data set with a different kernel configuration (the losing PO set). This is an instance of overhyping (overfitting due to selection of hyperparameters), because the hyperparameters determined with the PO set fitted the noise best compared to the other candidate hyperparameters. As the noise differed in the LB data set, classification performance was overall at a lower level and the pattern of more successful classification at later time points was also disrupted, causing our permutation test, introduced next, to generate significant AUC clusters for the winning PO, but not the LB map.

**Figure 2.**
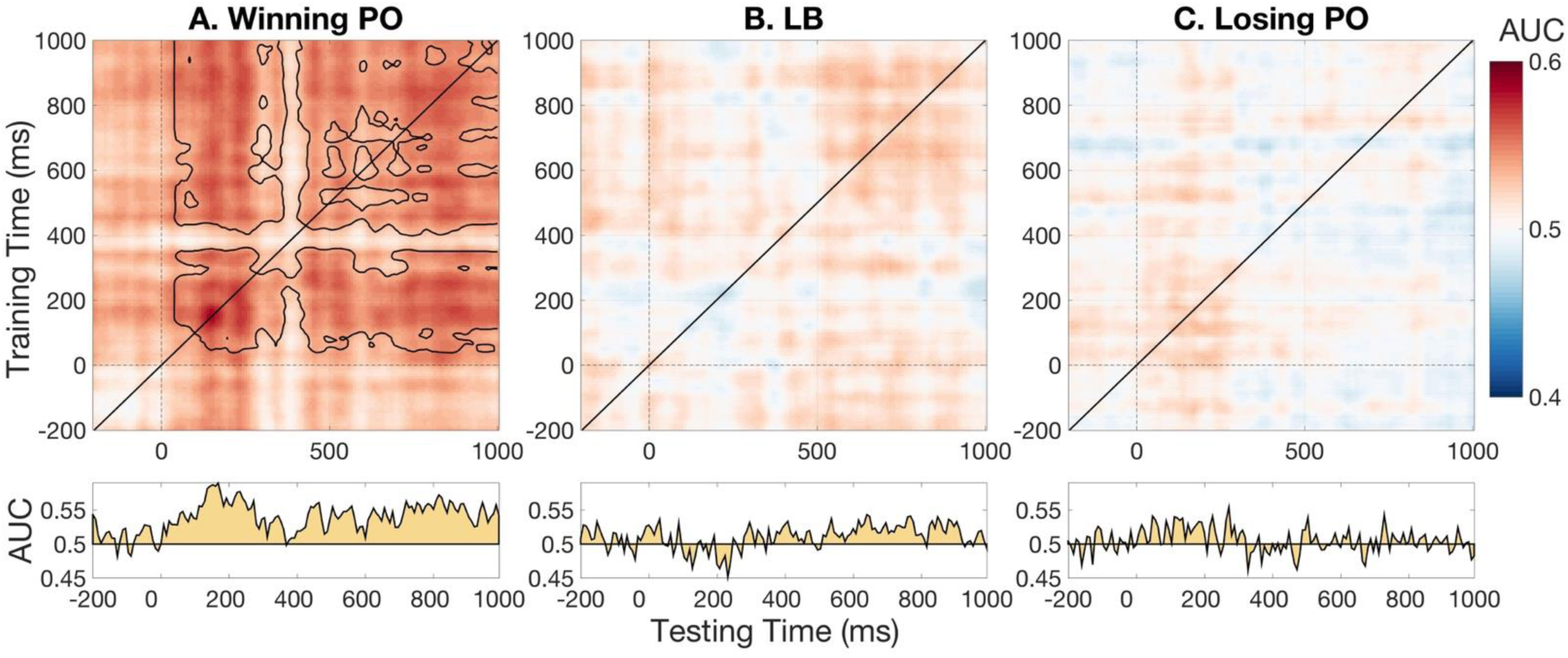
How overhyping manifests in temporal generalisation maps. Maps of a winning PO (panel A), its corresponding LB (panel B) and its worst PO, which implemented hyperparameters that led to *minimal* C-Mass, (panel C) are plotted with their main diagonal AUC vectors below. Beige areas in AUC time-series plots show divergence from chance-level classification (i.e. AUC of 0.5) in main diagonals. Classification performance was at a higher level for the winning PO compared to both other analyses. A family-wise error correction cluster-extent was performed for winning PO & LB maps and only showed statistically significant AUC clusters for the PO map. Maps and cluster-boundaries (i.e. matrices determining statistical significance) were 2D-smoothed separately using a boxcar of 40 ms width. This was only done to facilitate visualization and did not affect any analyses, which were all computed prior to smoothing. As all three analyses decoded null-data, any differences in classification performance must be due to the effectiveness of classifiers’ hyperparameters (in this case a linear SVM with a C parameter of 0.0059 for winning PO & LB). This is a demonstration of overhyping because these hyperparameters fitted the noise of the PO dataset best, which however differed in the LB dataset and thus led to decreased classification performance for the LB. This map triplet was manually chosen to show overhyping best. An additional 9 triplets can be found in the supplementary material.

We adopted a cluster-extent permutation test for our temporal generalisation maps, which was based on functions of the ADAM toolbox (J. Fahrenfort). We performed a first-level Wilcoxon signed rank test (non-parametric alternative to a t-test, preferred as distributions of AUCs do not meet parametric assumptions) at each pixel of the temporal generalisation map across single-subject maps, which resulted in a map of p-values. Neighbouring AUCs found to be significant for this test formed clusters and these clusters’ sizes are subsequently tested against a permutation-distribution of maximum cluster-sizes under the null. Clusters were determined statistically significant if only 5% of permuted maximum cluster-sizes exceeded their size (i.e. alpha of 0.05). For a more detailed introduction of this test, see Appendix 1.

To illustrate that these effects are systematic, a comparison of the PO competition winners against their respective LB counterparts shows how the C-Mass distributions are shifted by the selection process, despite the use of cross-validation (Figure 3). The top panel illustrates that the average C-Mass (which is always greater than zero since it is the sum of squares) is greater for the winning PO than the set of LB’s. The second panel illustrates that classification performance on a randomly selected set of hyper-parameters, as opposed to the winning set from the parameter-optimization phase, is approximately equal to performance on the LBs dataset when using the winning hyperparameter set, as it should be if the difference between PO and LB classification is due entirely to noise. This result demonstrates how a Lock Box provides an unbiased estimate of performance, as the resulting C-Mass is free of any overhyping effects.

**Figure 3.**
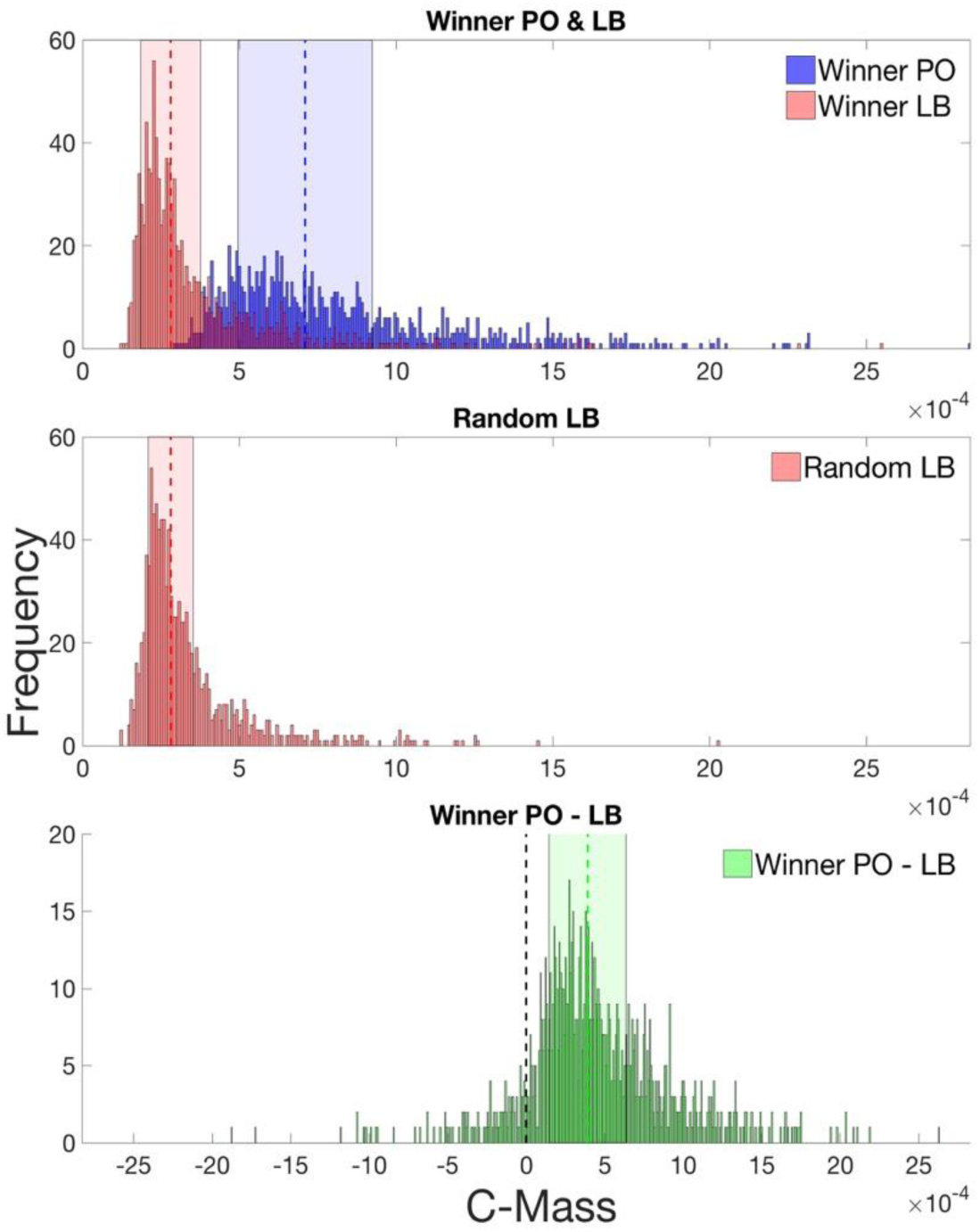
C-Mass distributions of PO (blue) and LB (red) maps (top two panels), as well as their within-iteration difference (bottom panel). The top panel shows C-Mass results for the PO & LB maps that incorporated winning hyperparameters from the parameter-optimization phase, the middle panel shows the distribution of LB C-Mass after choosing hyperparameters randomly. The coloured vertical lines indicate distributions’ median value and the rectangles surrounding these lines indicate the interquartile ranges. The black vertical line in the bottom panel indicates a PO – LB difference of zero (i.e. no overhyping). Note that even though LB C-Mass distributions correspond to simulated ‘null-distributions’, they are not centred around zero since we squared AUC values during the calculation of C-Mass.

The focus of this analysis lies in the top panel of Figure 3: the distributions of PO & LB C-Mass for winning hyperparameters of the PO competition clearly demonstrate overhyping of classification results. If overhyping was absent, these distributions should sit on top of one another. However, the PO C-Mass distribution has a higher mean (0.00081 / 8.1098*e-04), median (0.00071, 7.0974*e-04) and larger variance (1.3977*e-07) compared to the LB C-Mass distribution (mean: 0.000367 / 3.6650*e-04, median: 0.000281 / 2.811*e-04), variance: 6.217*e-08). The bottom panel of Figure 2illustrates how the *within-iteration* differences in C-Mass between PO and LB were distributed. This distribution should be centred around zero if no overhyping was observed (i.e. a given set of hyperparameters leading to similar success in decoding between-class differences for PO as well as LB null-data). The observed mean (0.000444 / 4.4448*e-04) and median (0.000391 / 3.9104*e-04) of the difference distribution was positive, implying higher C-Mass in PO maps. Permutation tests, which were based on randomly determining the direction of subtraction between PO & LB C-Mass to generate a distribution of null PO-LB C-Mass differences, confirmed that both values were significantly different from zero (p < 0.001), which provides evidence for overhyping. However, p-values obtained from simulation analyses should be interpreted with caution, as we discuss in more detail in the supplementary material.

We further assessed how vulnerable the different classifiers were to overhyping. Across all classifiers, the median difference in C-Mass between winning PO and LB was positive and significantly different from zero after performing the permutation test introduced above (linear SVM: median = 0.000353 / 3.533e-04, n = 299; polynomial SVM: median = 0.00036 / 3.58*e-04, n =176; RBF SVM: median = 0.00024 / 2.39*e-04, n =110; LDA: median = 0.000462 / 4.618*e-04, n =453). We investigated whether overhyping was more pronounced for certain classifiers by conducting a Kruskal-Wallis test (due to non-normality of C-Mass values), which revealed a significant difference among the four classifier types (χ^2^ (3, 1034) = 44.14, p < 2e^-7^). Post-hoc pair-wise tests of mean rank-differences between classifiers provided evidence that over-hyping was largest after LDA classification, smallest for RBF SVMs and intermediate for linear & polynomial SVMs (detailed results of this analysis can be found in the supplementary material).

Interestingly, we found that LDA classifiers won our PO competition most frequently. It was especially intriguing to see a clear trend towards hyperparameter settings which generate simple classification models (e.g. high regularization or a linear classification kernel) *winning and losing* our PO competition the most. Losing the competition means that a given setting of hyperparameters led to *minimal* C-Mass across all other settings in a given iteration. This rather counterintuitive finding can be explained by an analysis we present in the supplementary material, where we also display histograms of hyperparameter settings winning/losing our PO competition. We show with an additional set of simulations that C-Mass varies noticeably more after temporal generalization with simple classification algorithms. Such analyses hence yield more extreme values in both directions (low and high C-Mass) more often than complex classifiers, which results in simple classifiers being more likely to win *and* lose our PO competition. One could therefore argue that the risk of over-hyping due to not applying preventive measures such as a Lock Box is especially serious in the case of commonly used linear classifiers such as LDAs or linear SVMs.

### OVERHYPING BY FEATURE SELECTION DESPITE CROSS-VALIDATION

In addition to kernel parameters, analysis optimization can involve feature selection, in which portions of the data set are excluded from the pipeline on the grounds that they contain irrelevant information that can reduce classifier accuracy (e.g. Deshpande et al. 2010). This method is included as Recursive Feature Elimination (RFE) in scikit-learn. Here, we show that when cross-validation is the only protection against over-hyping, this method will induce spurious findings of significant classification accuracy on randomly shuffled data when feature selection is based on classification accuracy.

The raw data are the same as were used in the first analysis and are again randomly shuffled to remove differences between conditions (in a statistical sense). A simpler classifier is used which determines on each trial whether one or two targets had been presented based on the output of a spectral analysis. In this analysis, selection of features occurs by weighting different frequency components with channels collapsed.

## Methods

A fast Fourier Transform (FFT) of the 64 data points (comprising 256 ms) from each EEG channel after the first target onset were extracted from each trial. The log of the absolute value of the FFT was computed, and spectra across all channels were summed, resulting in 64 frequency values per trial. The classifier attempted to determine whether subjects had seen one or two targets within a given trial based on these 64 frequency values that represented the scalp-wide power spectrum from the 256ms time period after target onset. As above, the trial labels were randomly shuffled prior to the analysis to remove the correspondence between data and conditions.

A support vector machine (SVM) was used to classify the post-processed EEG data and the over-hyping was accomplished with a genetic algorithm that adjusted weights for the 64 frequency bands available to the classifier. The SVM was MATLAB’s fitcsvm, with an RBF kernel and kernelscale set at 25.

To demonstrate that cross-validation is inadequate protection against overhyping, the analysis was repeated for 16 iterations. For each iteration, 15% of the data were set aside in a Lock Box (LB) to test for over-hyping. Since the data was randomized, it was expected that performance on this outer test set should be at 50% (chance level), while performance on the 85% of the trials that formed the Parameter Optimization (PO) set would be elevated above chance by the last generation of the genetic algorithm. Over-hyping on the 16 PO sets was performed using a genetic algorithm coupled with cross-validation. For each generation of the genetic algorithm, 10 candidate feature-weight vectors were each evaluated against a shared set of 10 random partitions of the PO dataset, with 85% of trials in each partition used to train the SVM and 15% used for testing. At the start of the optimization procedure, the 10 candidate weight vectors were randomly constructed with 64 values ranging from 0.95 to 1.05. During training and testing, these vectors were multiplied by the power spectra for each trial before being provided to the SVM.

Within each PO iteration, for each of the 10 candidate feature weight vectors, the SVM performance in terms of AUC on the 10 random partitions was averaged to compute performance for each candidate. The best candidate was selected and then repeatedly mutated by adding 64 random numbers (range [-.05 .05]) to create 10 new candidates for the next generation of the genetic algorithm. This process was repeated for 400 generations to optimize the analysis.

To measure over-hyping, after each generation, the best feature weight vector was also used in a classification of the LB set for each of the 16 iterations and the results were not used to inform the evolution of the feature-weight vector. This is a strong violation of the principle of using a lock-box but it is done here as a demonstration. In practice accessing a lock-box multiple times can itself result in overfitting, particularly if the results are used to influence analysis choices or stopping criteria.

To measure the statistical significance of the model’s classification on the parameter optimization set, a permutation test was run after the final generation of the genetic algorithm. First, the analysis result was computed as the mean AUC across the ten PO partitions using the final generation of feature-weights. Then, all condition labels for the trials (i.e. the target-type) were randomly shuffled 1000 times. After each such shuffling, for each of the ten partitions, the SVM classifier was retrained with the best final weight vector and the AUC was measured. These AUC values were shuffled to create a null-hypothesis distribution of 1000 values. The p-value was then computed as the fraction of the null-hypothesis distribution that was larger than the non-permutated classification result (i.e. the proportion of shufflings that produced a mean AUC greater than the mean AUC on the unshuffled data).

This entire procedure was repeated independently for 16 iterations times to demonstrate the robustness of over-hyping. In each case, the data were randomly repartitioned into a parameter optimization set and a lock-box, and the genetic algorithm was used to optimize weights for the parameter optimization..

## RESULTS

The results of overhyping by feature selection are illustrated in Figure 4, which shows that performance improves on the parameter optimization set without corresponding changes on the lock-box set.. As the labels were randomly shuffled, any performance above chance (AUC of 0.5) in a statistical sense, would indicate overhyping. All 16 iterations of the PO set had significantly elevated performance by the final generation of the feature-selection genetic algorithm. One of the LB sets was significant.

**Figure 4.**
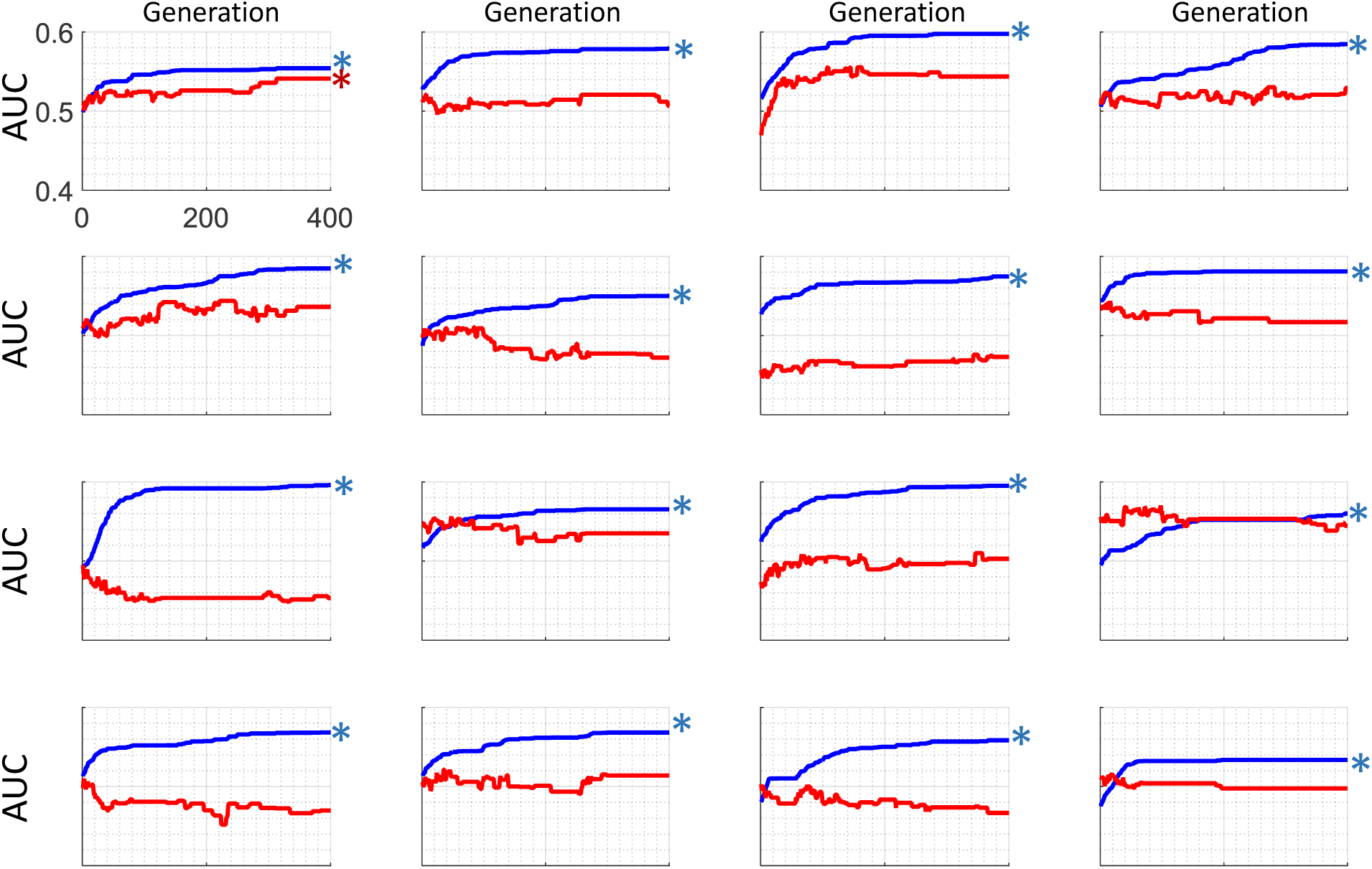
To demonstrate that models can be over-hyped using feature selection, a genetic algorithm was used to iteratively select features to optimize performance on a randomly shuffled EEG data set, thus performance should not deviate from chance. The optimization procedure was run for 16 iterations, with 400 generations in each. The blue trace indicates accuracy from a cross-validation test on the parameter optimization set, while the red shows performance on a lockbox set. The asterisks indicate when the results of the final generation differed significantly from chance at an alpha level of .05. All of the PO sets were significantly different from chance, while only one of the LB sets was.

## DISCUSSION

This paper demonstrates the ease with which over-hyping can be induced when using machine learning algorithms despite the use of cross-validation. The approaches used here are analogous to optimization procedures that have been used in EEG/MEG classification such as exploring various kernel options or discarding channels and frequency bands to improve classification performance. Similar problems may exist with other hyper-parameters, e.g. choosing time windows, or different ways of filtering out artifacts. Moreover, the same concerns apply to any kind of large neural data set. For example, in the case of using multi-voxel pattern-analysis (MVPA) on fMRI data, optimization through selection of any analysis step in the pipeline during consultation with the data could lead to the same kinds of over-hyping that we demonstrate here.

These results should not be taken to indict cross-validation as a poor methodological choice: It is considered to be state-of-the-art by many in the machine vision and machine learning communities for good theoretical and practical reasons (Arlot & Celisse 2010). However, our result does clearly indicate that cross-validation does not permit heedless analysis optimization.

There are several ways in which over-hyping can be protected against, above and beyond standard forms of cross-validation. We suggest that, in order to increase generalizability and replicability, journals publishing data from classification analyses encourage the use of one of the approaches listed below.

### THE LOCK-BOX APPROACH

Using a metaphorical data lock-box makes it possible to determine whether over-hyping has occurred. This entails setting aside an amount of data at the beginning of an analysis and not accessing that data until the analysis protocol is clearly defined, which includes all stages of pre-processing, artifact correction/rejection, channel or voxel selection, kernel parameter choice, and the selection of all other hyper-parameters. A close variation of this technique is already standard practice in machine learning competitions. When submitting a candidate for such a competition, the ultimate performance of the algorithm is evaluated on a separate set of data that is reserved until the final stage of the test. The workflow of using a lock-box is shown in Figure 5.

**Figure 5.**
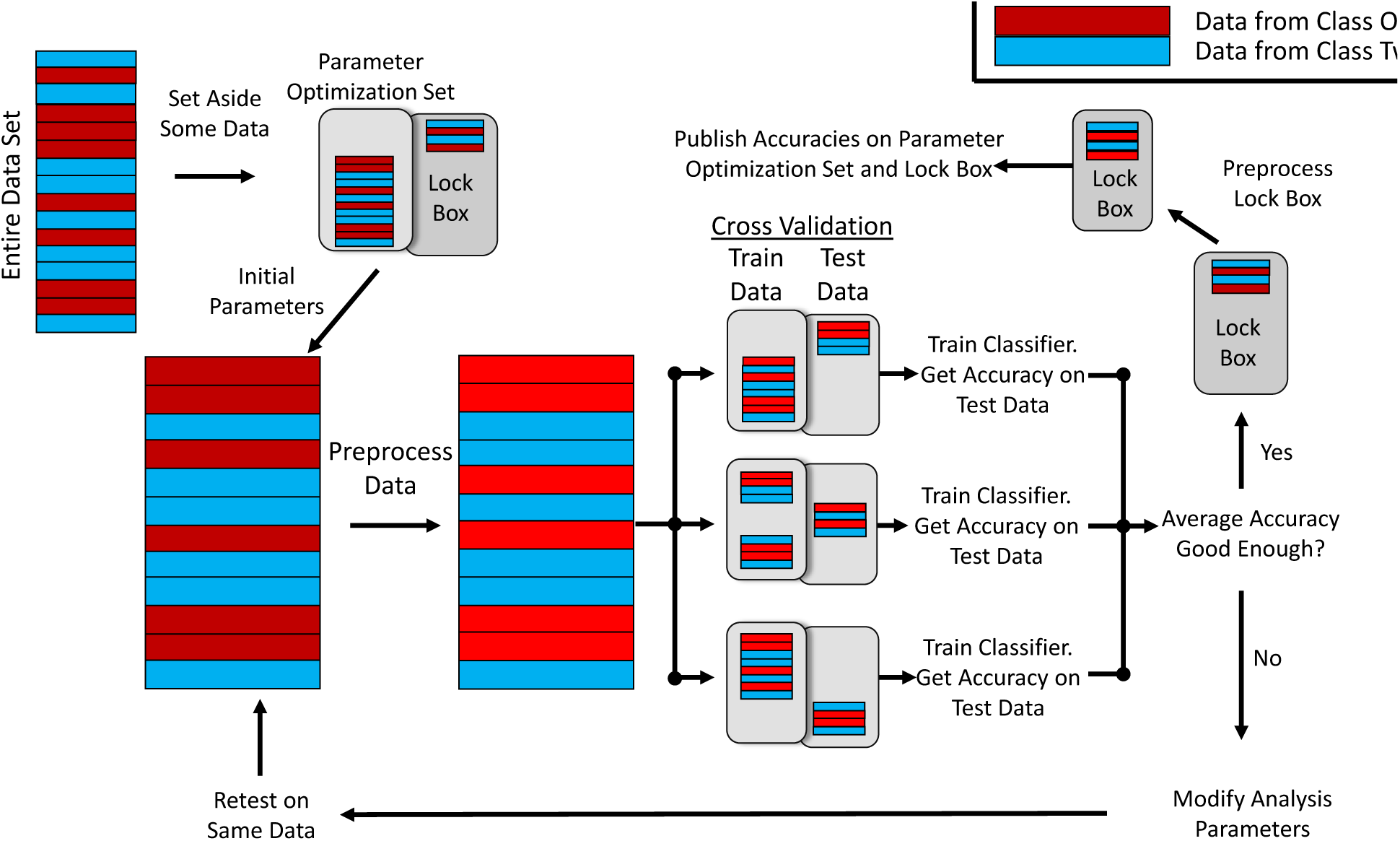
Here the workflow of using a lock-box is demonstrated in illustrative form. Data is first divided into a parameter optimization set and a lock-box. The model can be repeatedly tested and hyper-parameters can be iteratively modified on the parameter optimization set. After all hyper-parameter optimization and the analysis workflow is determined, the model can be tested against the lock-box data. By doing this, an unbiased estimate of overfitting can be obtained, and an objective measure of how well this system will generalize is achieved.

We suggest that, moving forward, machine classification approaches to data analysis in neuroscience routinely incorporate a lock-box approach, in which data are set aside at the beginning of the development of an analysis and not assessed until the paper is ready for submission (or equivalently, new data are collected at the end of analysis optimization). At this point, the data in the lock-box should be accessed just one time to generate an unbiased estimate of the algorithm’s performance. This result is likely to be less favorable than the data that were being used during optimization and should be published alongside the results from any other analyses.

If it turns out that the results from the lock-box test are unsatisfactory, a new analysis might be attempted, but if so, the lock-box should be re-loaded with new data, either collected from a new sample or from additional data that were set aside at the beginning of the analysis (but not from a repartitioning of the same data that had originally been used in the lock-box).

A possible alternative is to access the lock-box multiple times during optimization, but to apply a correction to any resultant statistics as a function of the number of times the lock-box data was evaluated. A method for accessing a lock-box multiple times while limiting overfitting was suggested by Dwork (2015). This method called for simultaneously evaluating a given model on both the parameter optimization set and on the lock-box, and then only revealing the performance on the lock-box to the operator if that performance was significantly different than that of the model on the parameter optimization set. Furthermore, the performance on the lock-box set would be presented only after being summed with a Laplacian noise variable. By following this method, the maximum error rate when generalizing to out-of-sample data can be limited by only observing the performance on the lock-box a set number of times (and halting parameter optimization once that limit is reached). While this is an innovative method for limiting overfitting, it only sets the maximum error rate when generalizing – To get the true error rate, a second lock-box would have to be used.

Note that this lock-box approach is evaluative. It does not prevent over-hyping, but allows one to test whether it has occurred. However, the performance of the algorithm on the lock-box is guaranteed to be a non-over-hyped result if the technique was correctly used.

### NESTED CROSS-VALIDATION

Another way to respond to the problem of overfitting hyper-parameters is to use a generalization of cross-validation, called *nested cross-validation* (Cawley & Talbot 2010; Stone 1974). Nested CV helps to ensure that results are not specific to a given analysis configuration by showing that the results generalize to out-of-sample data. In this approach, inner cross-validations are run within an outer cross-validation procedure, with a different portion of the data serving as outer “hold-out set” on each outer iteration. Importantly, for each outer iteration, an unbiased assessment of accuracy can be obtained by testing on this outer hold-out set. That is, the best parameters and hyper-parameters determined on each inner cross-validation, can be assessed out-of-sample on the corresponding outer hold-out set.

Nested cross-validation can be thought of as a repeated lock-box approach, in which a new box (the hold-out set) is set aside and locked for each iteration of the inner cross-validation loop (Figure 6). Then, an overall accuracy (and indeed dispersion of accuracies) can be obtained by averaging across the accuracies determined from the hold-out sets of each outer iteration. This will typically be a more reliable measure of accuracy than that obtained from any individual outer iteration (i.e. the lock-box approach). However, it is critical that the outer folds are not cherry-picked to find the best solutions, since this would constitute over-hyping. It is also important that the algorithm not be re-run in its entirety with different parameters after viewing the results, since this again would result in over-hyping.

**Figure 6.**
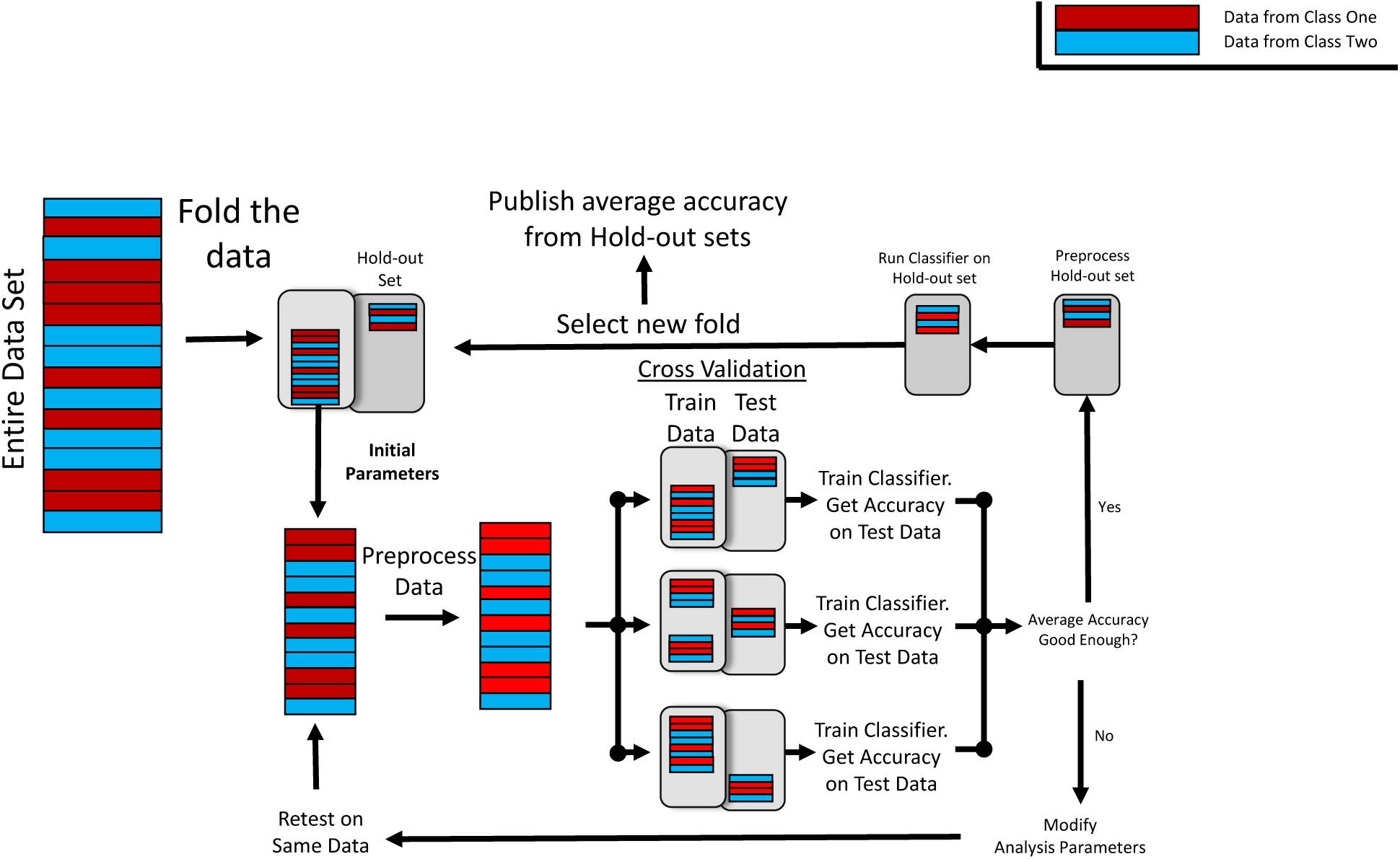
Here the workflow of nested cross-validation is demonstrated in illustrative form. The data set is folded into multiple combinations of hold-out set and inner optimization set. Each of these folds is essentially similar to the lock-box approach described above and can be optimized. The final accuracy would be the average accuracy computed across all of the hold-out sets.

An issue for nested cross-validation relative to the lockbox is that the average accuracy obtained at the end of the procedure will be the result of multiple configurations of hyper-parameters, and thus it may be especially difficult to understand the link between the data and the accuracy. For example, in analysis of fMRI data where the region of interest is one of the hyper-parameters, different iterations of the outer loop may converge on different regions of the brain. It would therefore be difficult to gain insight into what brain areas are driving the classification. We give more details of nested cross-validation in the Supplementary Material.

### THE BLIND ANALYSIS APPROACH

Blind analysis is an appropriate tool for preventing over-hyping when testing a well-defined hypothesis. Under a blind analysis scheme, the input data is modified in a reversible way to prevent the optimization process from inadvertently becoming tuned to noise patterns in the data that are differential with regard to the contrast of interest. An alternative is to use an orthogonal contrast, where classification is done on a condition that is orthogonal to the classification one will ultimately use (Brooks et al. 2017) Some examples of using blind analysis include scrambling all condition labels and then artificially adding ‘target signals’ to some trials. The hyper-parameters of the model can then be optimized to detect the signal present in the modified data. Once the hyper-parameters are locked in, the blind can be lifted (e.g. conditions unscrambled and modifications removed), and the true results can be calculated. The advantage of this approach is that all of the data can be used during the optimization phase, and the final evaluation of performance can be done across all of the data instead of just the outer-box set. Note that blind-analysis is a way to minimize over-hyping. If used in conjunction with a lock-box, one can both minimize and diagnose overfitting. The disadvantage of the blind analysis is that it obscures accuracy on the key predicted variable, and this may prevent the development of an effective analysis plan depending on the type of data one uses, in which case a lock-box is a good solution.

### THE PRE-REGISTRATION APPROACH

Cross-validation does succeed in providing an unbiased estimate of out-of-sample performance when classification results are not used to iteratively optimize performance. Therefore, it is safe to pre-register or otherwise rigidly specify a classification analysis before attempting it. Assuming that it is done in good faith, the pre-registration would provide evidence that the hyper-parameters were finalized prior to attempting the analysis using previously established methods. The advantage of this approach over the lock-box is that all of the data can be used in the final estimate of performance. The disadvantage is that hyper-parameter optimization is not permitted, which limits the effectiveness of the analysis.

### WHICH APPROACH TO USE

Each of these approaches is ideal for particular use cases. The simplest decision point hinges on whether the analysis plan is already established, in which case pre-registration is clearly the best choice. Blind analysis is suitable when hyper-parameters need tuning to accommodate unanticipated variability in the data that is orthogonal to the predictor (e.g. finding the time window or location of a brain signal of interest). Nested cross-validation is well suited to a case in which an automated algorithm can be used to tune hyper-parameters, and the precise values of those hyper-parameters are not of interest. Finally, the Lock box, particularly when it is very large, is best suited to a case in which the values of tunable hyper-parameters are of particular interest or the process of tuning them is done partially by hand, rather than by automation.

Regardless of which approach one takes, it seems crucial that more transparency should be applied to documenting how data is treated through the entire process of developing a pipeline. For example pilot tests of an analysis can lead to overhyping if they inform the search range of hyper-parameter optimization prior to partitioning data into different sets. In such cases, being transparent can highlight the points where leakage of information into the (hyper-parameter dependent) pipeline may have occurred.

### SAFE VERSUS EFFECTIVE USE OF MACHINE LEARNING

Optimal use of machine learning in neuroscience requires that it be used both safely (i.e. without over-hyping such that the results can be trusted) and effectively (i.e. the classifier is appropriately tuned to discriminating signal). In the terminology of machine learning, *safe* largely corresponds to *minimizing variance*, while *effective* largely corresponds to *reducing bias* Geman, Bienenstock, & Doursat (1992). The methods described above help to ensure safety, but do not necessarily provide effective solution, since the avoidance of over-hyping is often obtained by limiting the amount of analysis optimization that is allowed. When data is easy to obtain, this limitation is not as severe, since analysis chains can be repeatedly adjusted, and tested against new data. However data in the neurosciences is often expensive and time consuming to collect. Unfortunately, this means that one often has to choose between analyses that are highly optimized but over-hyped, or weakly optimized and not over-hyped. The best path forward is to make use of expertise when it is available, such that good decisions are made up front, and ideally even pre-registered prior to viewing the results of analysis on critical data.

## CONCLUSION

The biggest danger of data science is that the methods are powerful enough to find apparent signal in noise, and thus the likelihood of data over-hyping is substantial. Furthermore, analysis pipelines are complex, which makes it difficult to clearly understand the possibilities for leakage between optimization and evaluation stages that can lead to over-hyping. Our results illustrate how easily this can occur despite the use of cross-validation. Moreover, it can be difficult to detect over-hyping without having an abundance of data, which can be costly to collect. However, as reproducibility is a cornerstone of scientific research, it is vital that methods of assessing and assuring generalizability be used. By setting aside an amount of data that is not accessed until the absolute completion of model modification (i.e. a lock-box), one can obtain an unbiased estimate of the generalizability of one’s system and of how much over-hyping has occurred. Alternatively, blind analysis methods, good faith pre-registrations of the analysis parameters and nested cross-validation reduce the possibility of overfitting. Conversely, using any method that allows one to check performance on the same data repeatedly without independent data that has not been consulted can induce over-hyping, inflating false positive rates and damaging replicability. Devoting more attention to these dangers at this point, when machine learning approaches in neuroscience are relatively nascent, will allow us to improve the state of science before inappropriate methods become standardized.

## Supplementary Material

We present more detailed descriptions of cross-validation procedures.

### Cross-validation

Cross-validation procedures are defined according to the number of folds they employ. For example, in 10 fold cross-validation, the data are randomly divided into 10 equal sized portions, and a 10 iteration procedure is performed. In each iteration, a different one of the 10 portions is used as the test set, with the remaining 9 unioned together to provide a training set. An accuracy is thereby obtained for each iteration, which critically, is “out-of-sample” with respect to that iteration, since the test set is disjoint from the training set. The results of each iteration (here 10) are then averaged, to provide an overall accuracy. The procedure has the advantage that even though an out-of-sample test is performed, all data points appear at one instant in a test set, ensuring that an accurate assessment of overall classification accuracy (which is averaged across all iterations) is obtained. Additionally, an assessment of the confidence one should have in the overall accuracy can be obtained from the dispersion of the accuracies generated across cross-validation iterations. As demonstrated in this paper, a single execution of cross-validation is an effective and safe way to fit parameters, but if the entire procedure is run multiple times to fit hyper-parameters, it has the potential to overfit.

### Nested Cross-validation

This is a procedure in which cross-validations are nested within cross-validations. A typical approach would be to nest a cross-validation that provides an out-of-sample test of the fitting of parameters, inside a cross-validation that provides an out-of-sample test of the fitting of hyper-parameters. Critically, the procedure enables out-of-sample tests at both levels, i.e. inner test sets are out-of-sample re. the setting of parameters, while the outer test/hold-out sets are out-of-sample re. the choice of hyper-parameters. Effectively, the outer-level test/hold-out set really acts like a lock-box, but specific to a particular outer iteration. In this way, each data point contributes to a hold-out set at some instance. Accordingly, one obtains a better estimate of overall out-of-sample accuracy, since multiple hold-out set accuracies are calculated and averaged, rather than a single one, as would be the case with a single (fixed) lock-box and one level of cross-validation. Additionally, one obtains a measure of confidence of hold-out accuracies by considering the dispersion of such accuracies across the outer folds of the procedure. It is critical that accuracy from all of the outer loops is averaged together however. One cannot select the most favorable result without overhyping.

Figure S1 presents a typical nested cross-validation scheme. Importantly, as just discussed, with such a procedure, one can fit hyper-parameters as well as parameters. For example, if there were four Support Vector Machine kernels that one wanted to select between, one would run each of the three outer iterations shown in figure S1 four times, e.g. outer 1, 4 times (giving 4 × 2 = 8 runs for that outer iteration). One would then have an inner cross-validation accuracy for each kernel on each outer iteration, i.e. 4 for each outer iteration, with each of these 4 being obtained by averaging across the accuracies obtained on the corresponding two inner iterations.

One can then take the best kernel of the 4 tested, with its trained parameters, and test it out-of-sample on the hold-out set for that outer iteration. The hold-out set accuracies obtained from each outer iteration can then be averaged, to give a fair accuracy for the entire fitting – choice of kernel and fitting under each kernel. However, selectively reporting the best of those four kernels and ignoring the other three would constitute overhyping.

**Figure S1:**
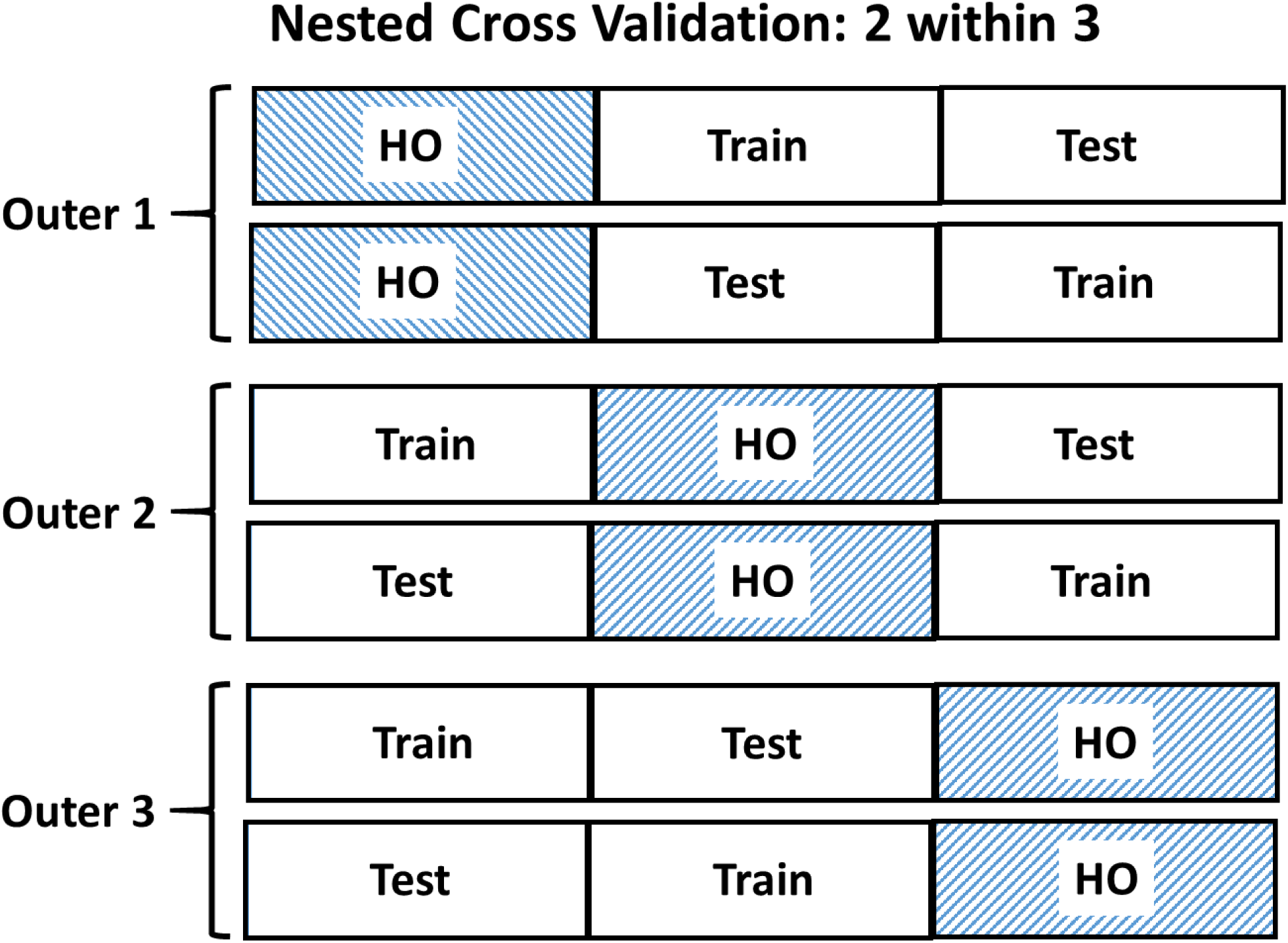
A 2 within 3 Nested Cross-validation, where the outer cross-validation loop involves a different Hold-Out (HO) set for each of its three iterations. The remainder of the data provides a parameter optimization set for each outer iteration, which is subjected to two fold (i.e. split-half) cross-validation.

### Cluster-Extent Permutation Test of Temporal Generalisation Maps

To family-wise error (FWE) correct our temporal generalization maps we performed the following procedure. For each group map, we first subtracted chance-level AUC (i.e. 0.5) from all its underlying single-subject temporal generalisation maps. Each position (pixel) of the single-subject maps was then tested separately against a median of zero across subjects using a two-tailed Wilcoxon signed-rank test. This yielded a map of p-values, which was subsequently used to form clusters if (at least 8-pixel) neighboring AUCs had p-values smaller than or equal to 0.05 and were of the same polarity. Polarity in this context means that we differentiated between positive (above-chance) and negative (below-chance) clusters. The unlikely case of adjacent & significant above- and below-chance AUCs would hence not form the same cluster. We recorded the sizes of these clusters and proceeded to computing the permutations.

We computed a total of 1000 permutations for this test. For each permutation, we performed a sign-swap at the level of single-subject maps to simulate the null. Specifically, instead of subtracting chance-level AUC (0.5) from *all* single-subject maps, as described above, we now randomly determine, with equal likelihoods and *for each single-subject map separately* whether 0.5 was subtracted from it (map – 0.5) or whether *the whole map was subtracted from 0.5* (0.5 – map). We again perform two-tailed Wilcoxon signed-rank tests at each pixel of our permuted single-subject maps, get maps of p-values and form clusters, as described before. After recording all clusters’ sizes, we determine the biggest cluster across both polarities (above- and below-chance) and store its size in a distribution. Repeating this process 1000 times results in a distribution of 1000 maximum cluster-sizes under the null against which our observed clusters were tested. Observed clusters were assigned p-values as the proportion of maximum (or biggest) clusters under the null found to be equal to or larger than them.

This procedure was always performed separately for each group map tested (meaning permuted distributions of maximum cluster-sizes were never used more than once). We also restricted the tested area to be from approximately 100 milliseconds onwards to resemble real-life usage of this test. It should be noted that this does not change the statistics in any way, because tests of maximum statistics are automatically correcting for family-wise errors (FWE) as long as one is adopting the same approach (e.g. restricting the area) for the observed and the (permuted) null data (in this case temporal generalisation maps).

### Degree of over-hyping across different classifiers assessed with Kruskal-Wallis Test

The Kruskal-wallis test is a non-parametric alternative to a one-way ANOVA, which we used for assessing the degree of over-hyping across different classifiers because our C-Mass distributions did not meet parametric assumptions. It was computed on classifier-separate C-Mass difference distributions between PO and LB (compare bottom panel of Figure 3). It hence tested the null hypothesis that the PO – LB C-Mass differences of our four groups (i.e. classifiers) originated from the same distribution. This null hypothesis was rejected (χ^2^ (3, 1034) = 44.14, p < 2e^-7^), as can be seen in the ANOVA Table below (Figure S2, panel A). Following up on this result, we further tested pair-wise mean-rank differences between classifiers using Tukey’s honest significant difference criterion to correct for multiple comparisons. The results of these tests are provided in Figure S2 panels B & C and showed that: 1) LDA had a significantly higher mean rank than all other classifiers (i.e. most over-hyping); 2) RBF SVM had a significantly lower mean rank than all other classifiers (i.e. least over-hyping) and 3) linear as well as polynomial SVM had moderate mean ranks, which were significantly smaller than LDA’s and larger than RBF’s but not significantly different from each other. These results thus provide evidence that the degree of over-hyping can vary substantially across classifiers.

**Figure S2:**
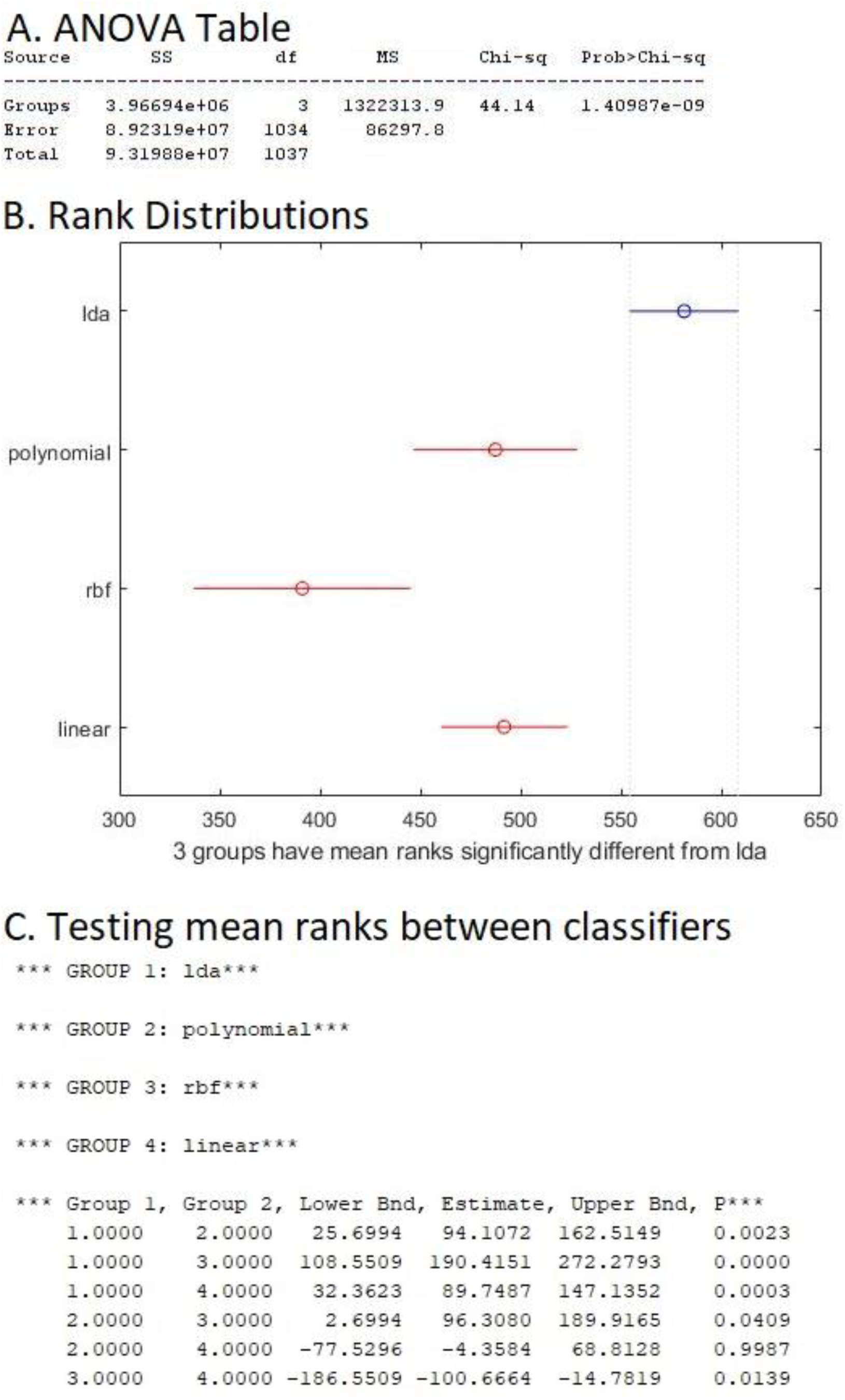
Results of Kruskal-Wallis test conducted on PO – LB C-Mass distributions to assess the degree of over-hyping between our four classifiers. Panel A shows the ANOVA table, which indicates a significant main effect of classifier. Panel B plots the rank distributions for each classifier, which indicates that mean ranks (i.e. degree of over-hyping) of LDA > linear / polynomial SVMs > RBF SVM. This is confirmed in Panel C, which shows results of statistically testing all pair-wise mean-rank differences. Five out of six tests yielded significant p-values after correcting for multiple comparisons using Tukey’s honest significant difference criterion. Only the contrast of linear vs. polynomial SVMs had a statistically non-significant mean-rank difference.

### Interpreting p-values in the context of simulations

P-values can become uninformative in the context of simulations, where very large simulated samples can be run. As discussed in Friston (2012), the fallacy of classical inference states that once the sample size is sufficiently large, p-values become trivial as the smallest effects suffice for significance. To be more precise, for a two-sided test, there is a sample size N for any non-zero experimental effect, no matter how small, that will make the p-value significant. We nonetheless chose a large sample size because, in contrast to p-values, standardised measures of effect size become more accurate with increasing sample size. This was recently illustrated in a neuropsychology context (Lorca-Puls et al., 2018, especially figures 4 & 5).

### C-Mass variance differences across different classifiers

We ran an additional set of simulations for this analysis, which was very similar to the one introduced in the “Overhyping due to kernel selection despite cross-validation” section of the main paper. We used the same dataset and shuffled class-labels randomly to simulate null-data. However, we did not group subjects into two groups of Parameter Optimization and Lock Box, but instead ran all combinations of hyperparameters (the same ones as introduced in the paper) in each of 1000 iterations on the data of all 24 subjects. To reiterate, this involved, for each iteration and configuration of hyperparameters, running a temporal generalization analysis on a given subject’s (null) dataset and then averaging all 24 single-subject maps to generate group-level maps. This therefore resulted in a total of 40000 group-level temporal generalization maps (1000 iterations * 40 hyperparameter configurations). We computed C-Mass as the average deflection from chance-level classification of a given group-map, as explained in detail in the paper. Figure S4A displays boxplots of these 40000 C-Mass values, which illustrates that C-Mass varies more after adopting simple classification algorithms (e.g. due to a high degree of regularization), even though the median C-Mass value is comparable across hyperparameter settings. Note that the degree of regularization decreases from left to right and models hence become increasingly complex. This means that temporal generalization maps were more likely to be either closer to chance-level accuracy (low C-Mass) or further away from it (high C-Mass) after running more simple classification algorithms, as such algorithms are more likely to yield *more extreme* C-Mass values. We further present in Figure S4B how this dynamic manifested in our PO competition, where the simpler classification models were, the more often they *won and lost* (the latter implying *minimal* C-Mass within a given iteration across all other hyperparameter settings). This analysis overall suggests that over-hyping due to the absence of appropriate precautions such as a Lock Box or nested cross-validation is more dangerous when adopting more simple classification models, such as the commonly used LDA and linear SVM classifiers.

**Figure S3:**
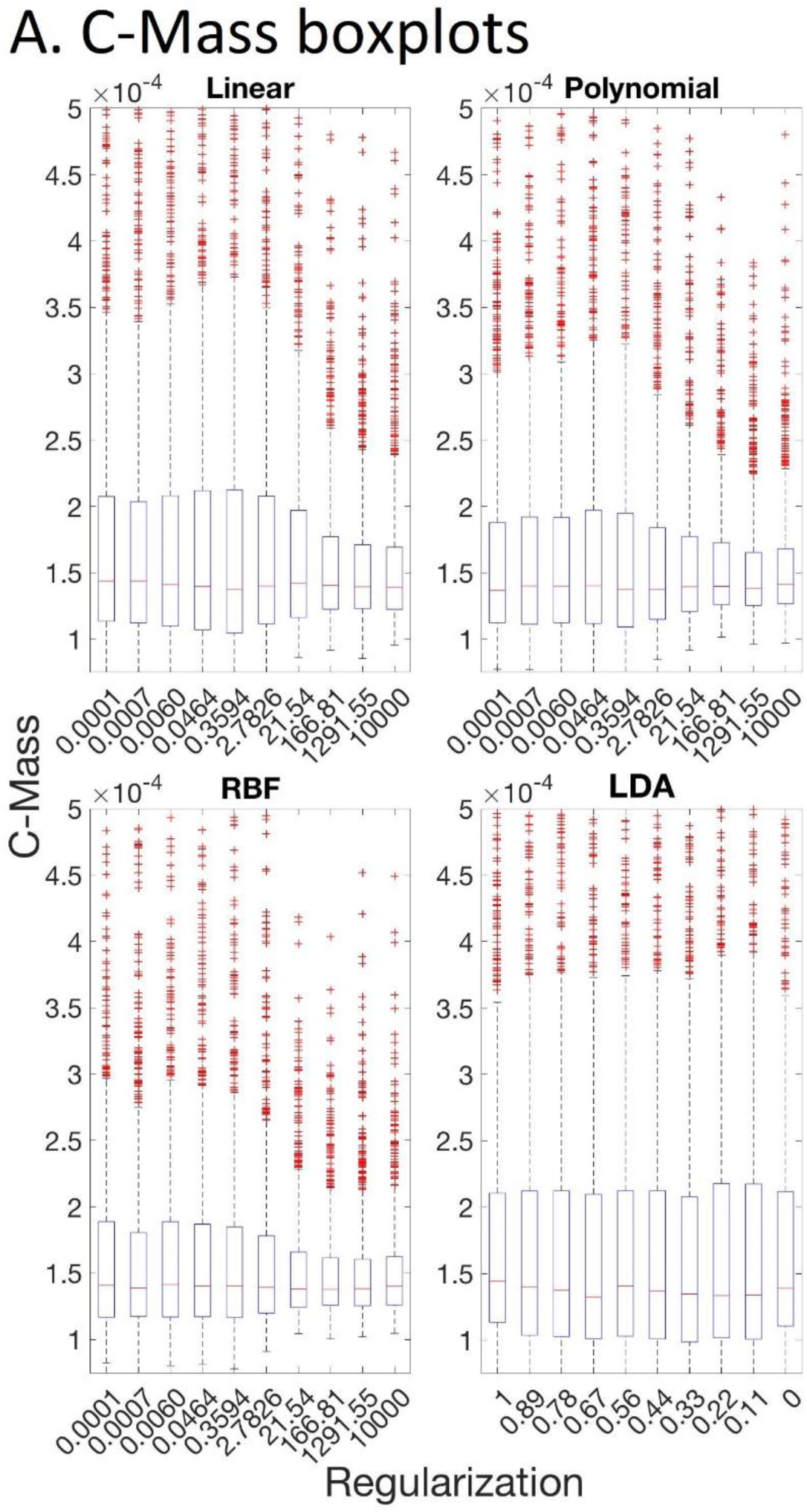
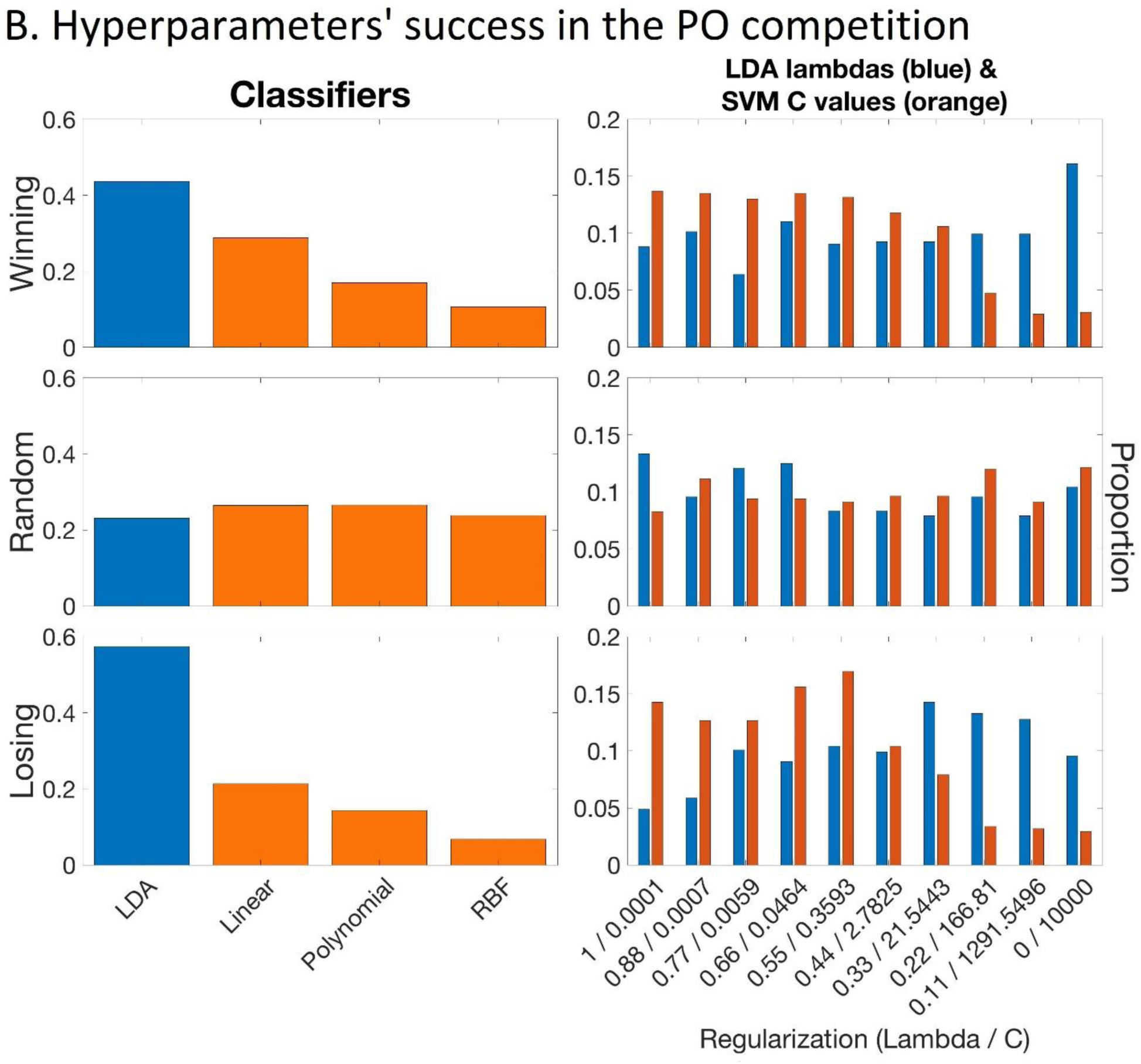
Panel A shows boxplots of C-Mass for all hyperparameter configurations. Red horizontal lines indicate the medians. Note that this analysis did not involve splitting subjects into PO and LB groups. Panel B displays proportions of hyperparameters selected after winning (top row) or losing (bottom row) the PO competition. The middle row corresponds to randomly selected hyperparameters, which were applied to the LB independent of the PO competition. The left column depicts distributions of classification models (blue = LDA, orange = different SVMs) and the right column shows how regularization parameters (lambda (blue) for LDA and C (orange) for SVM classifiers) were distributed. Note that regularization decreases from left to right in both panels (classification models become increasingly complex).

### Nine additional and randomly chosen temporal generalization plots for further illustration of over-hyping

Figure 2 of the main body illustrates just one of the 1038 results that was obtained. Here we show 9 additional samples from this distribution of 1038. These were selected at random, with bias neither towards significance of clusters, nor C-MASS. Five of the nine simulations selected at random have significant clusters in the Winning PO.

**Figure S4:**
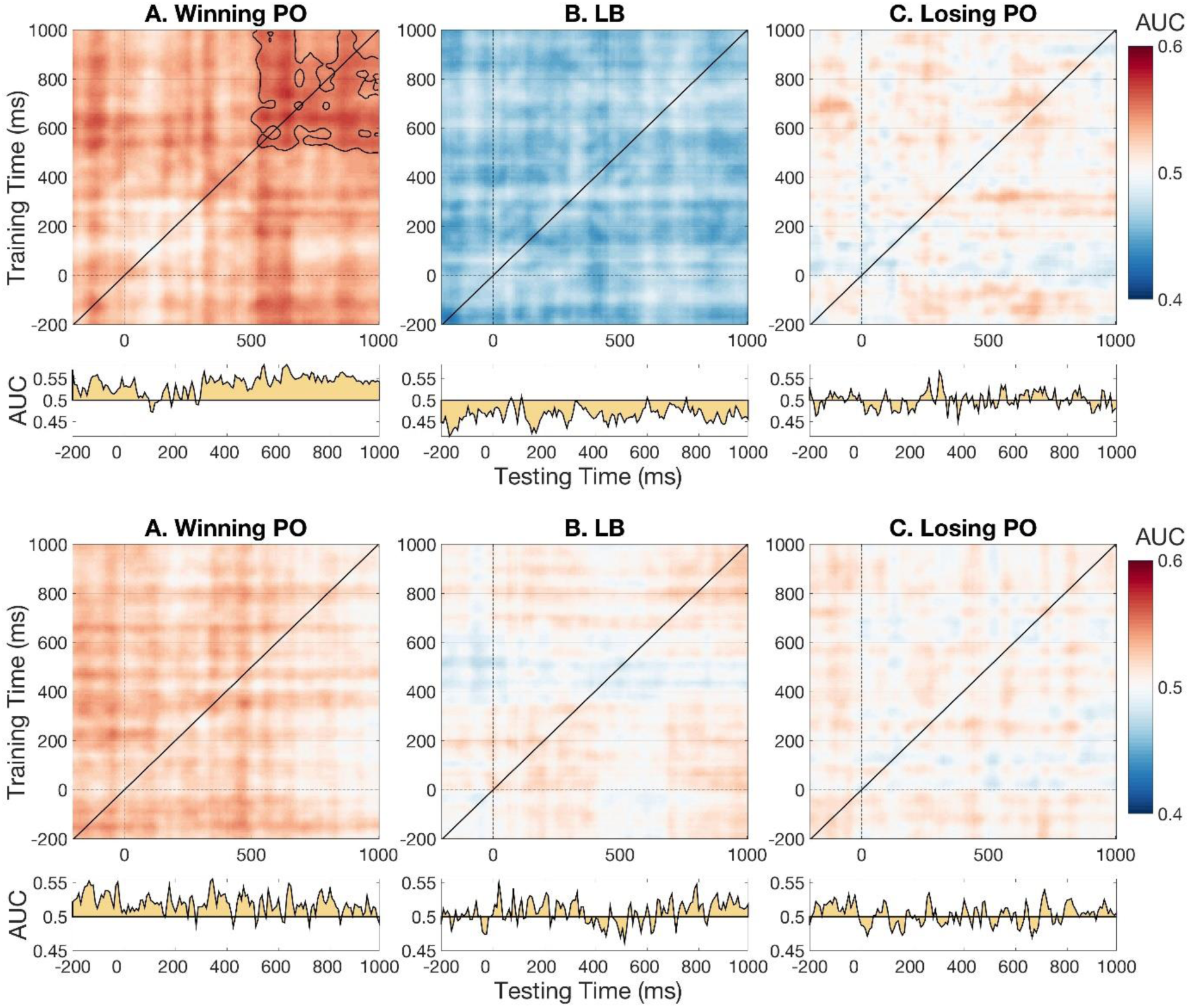
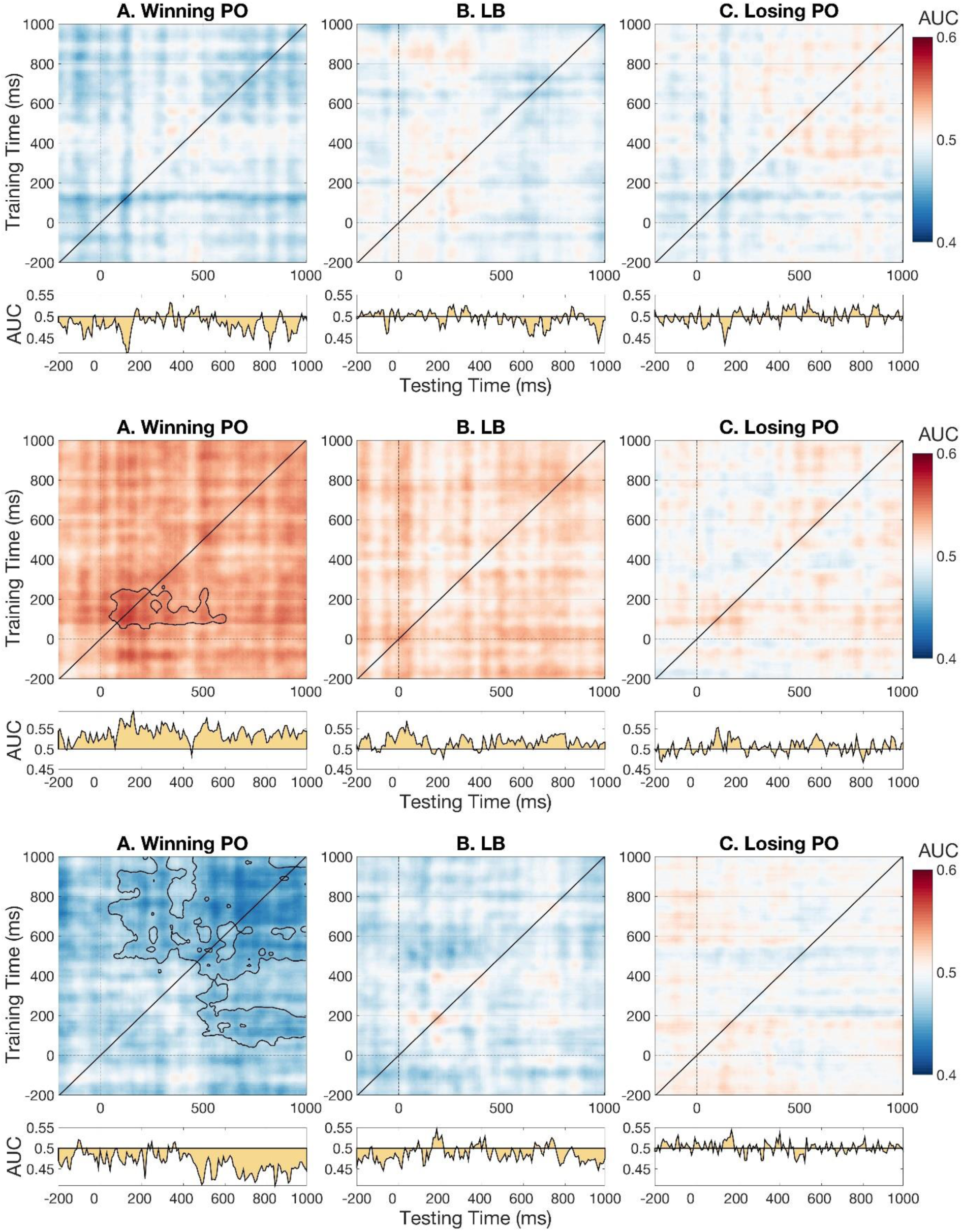
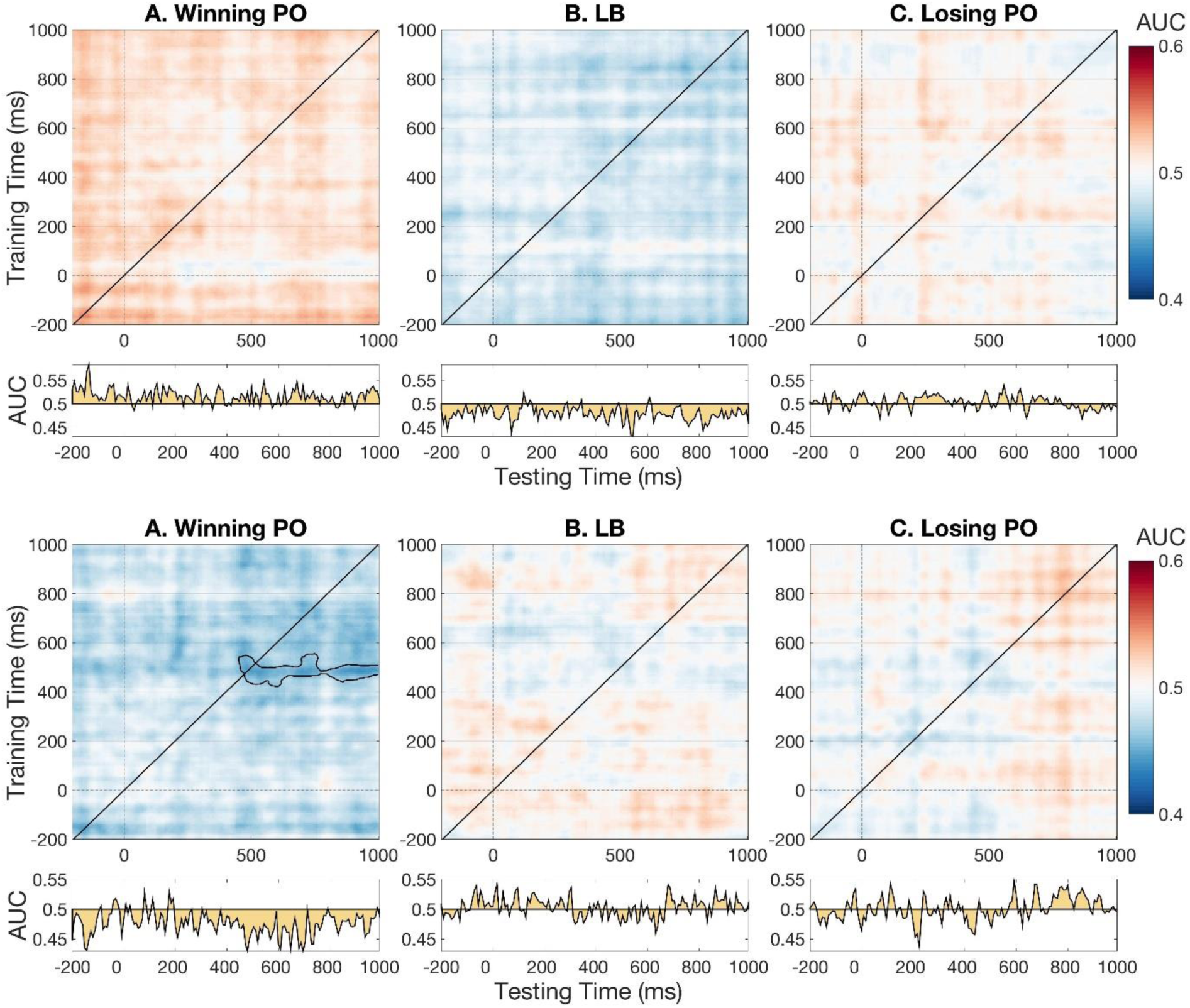
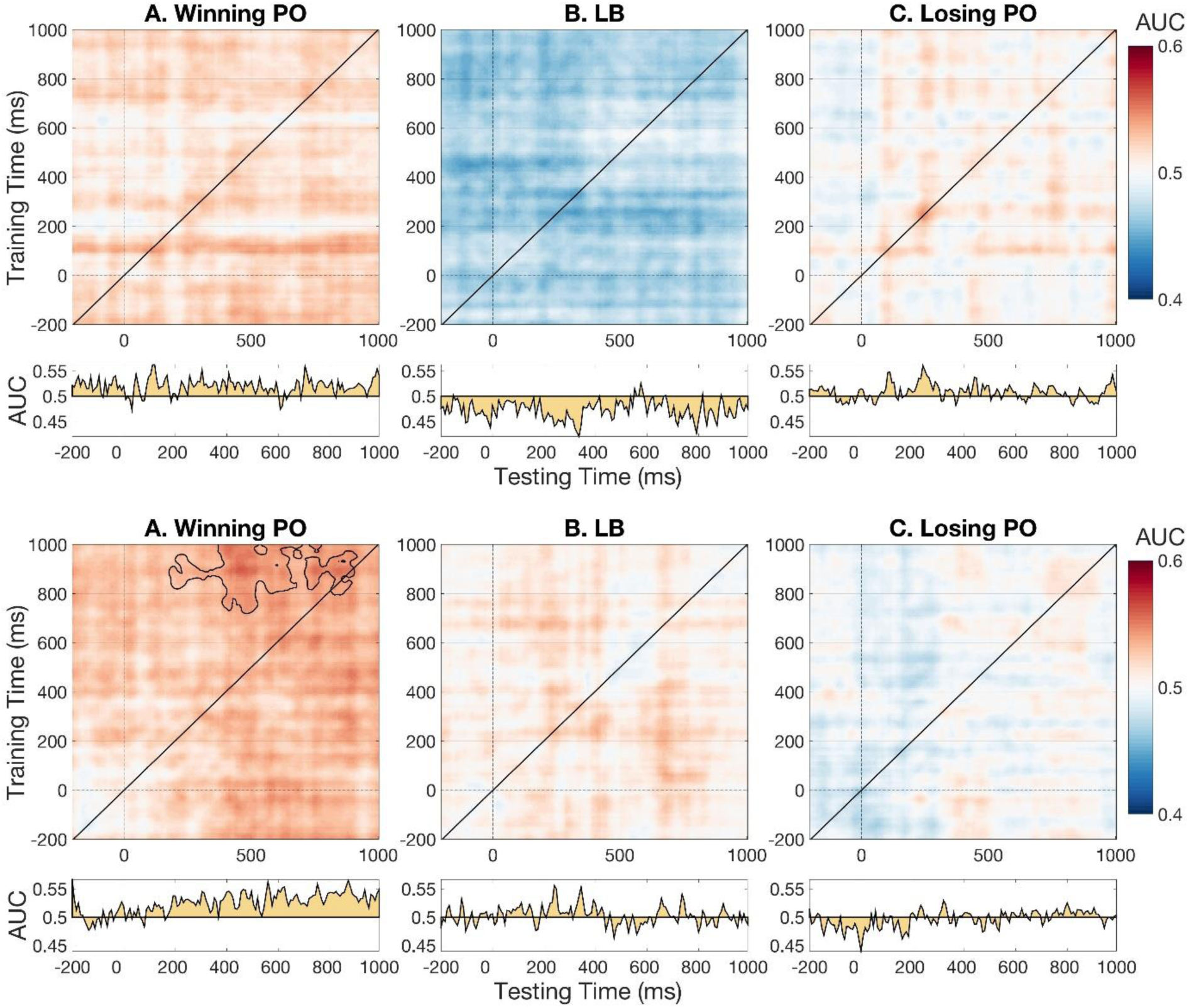
Nine additional temporal generalization map-triplets showing over-hyping as demonstrated in Figure 2. These nine candidates were selected randomly and correspond to iteration numbers 129, 994, 1002, 231, 164, 1008, 504, 83 and 846 (in the order plotted above) of our PO competition. Panels A, B and C show the winning PO, the (winning) LB and the losing PO maps, respectively. Plotting conventions follow those of Figure 2. Temporal generalization maps and permutation tests’ cluster-boundaries were again smoothed only to improve visualization and did not affect any analysis whatsoever.

### Methods from Callahan-Flintoft et al. 2018, Experiment 3

These are the methods used for collecting the EEG data set from Experiment 3 of Callahan-Flintoft et al 2018. These data were used in all of the analysis here.

#### Participants

Data from 26 subjects were collected for the first experiment. All subjects volunteered from the Pennsylvania State University psychology subject pool for this study. Participants all had normal or corrected-to-normal vision and were between the ages of 18 and 23 years old. Informed consent was obtained for each participant prior to each study in accordance with the IRB office of Penn State. Subjects were excluded for having too few usable trials per condition (threshold: 15) after discarding trials for inaccurate responses and EEG artifacts, leaving 25 subjects usable for analysis.

#### Stimuli & Apparatus

Participants sat in an electrically shielded room, 91cm away from the computer which had a 46cm CRT (1024 × 768, 60 Hz refresh rate). Stimuli were black (RGB([0,0,0]) alphanumeric characters in size 55 Arial font (1.26° × 0.63° of visual angle), presented on a grey background using MATLAB 2012 with Psychophysics Toolbox-3 extension (Brainard, 1997). Subjects looked for digit targets presented among a bilateral stream letters. The digits used as orthographic targets were: 2,3,4,5,6,7,8,9. Distractors and feature targets were drawn from the following collection of letters: A,B,C,D,E,F,G,H,J,K,L,N,P,Q,R,T,U,V,X,Y. Letters that resembled digits, such as ‘O’, ‘S’, ‘Z’ and ‘I’, were excluded as well as wider letters such as ‘M’ and ‘W’. Target identities were randomly selected on each trial.

#### Procedure

On every trial participants were presented with two RSVP streams placed three degrees from a fixation cross, which remained throughout the trial (Figure 5). Both streams were updated simultaneously with a 150ms stimulus onset asynchrony (SOA) and no inter-stimulus-interval. Note that this SOA is slightly longer than that used in Tan & Wyble (2015). This change was implemented to increase accuracy, since analyses could only be done on trials in which subjects reported both targets successfully. Seven or eight distractors in each stream were presented before target onset and eight distractor pairs were presented afterwards. This number of pre- and post-target distractors was used in all experiments except Experiment 2. Every trial had one or two targets, the second of which (T2) always appeared in the same stream at a varying lag. No stimulus was repeated for at least two sequential frames. At the end of the trial, the fixation cross was replaced with either a period or a comma for 150ms. Two practice trials were used within the instruction block to demonstrate the task.

Four conditions were used to test what duration between targets is required to elicit a second N2pc. Participants were either presented with only one target (Single), two sequential targets, the second immediately after the first (Lag 1), two targets with one frame of distractors in between (Lag 2), or two targets with three frames of distractors in between (Lag 4). There were 280 trials distributed equally across the four conditions, intermixed within a single block. After every 20 trials, participants were given a self-paced break.

##### Instructions and responses

Participants were told that every trial would contain one or two targets that they would have to report, in the correct order, using the keyboard. They were also told that on some trials they would have to report which symbol had replaced the fixation cross. At the end of each trial, participants were asked for the first and second target, with the option of pressing Enter if they did not know it. On a third of trials participants were also asked to report the symbol (dot or comma) that appeared at fixation. This was done in order to encourage participants to remain engaged, with their eyes on fixation, for the entire trial and not just until the target(s) appeared. This technique was used to discourage eye movements but the actual elimination of trials with eye movements was done with EEG measures described below. Feedback was provided by showing the participants what targets in order were shown as well as what punctuation had replaced the fixation cross (dot or comma).

Participants were excluded from both behavioral and EEG analysis if they had fewer than 15 usable trials in any condition. Trials were considered usable if participants accurately reported the identity of all targets present, regardless of the order, and if the trial passed the EEG exclusion criteria described below. The average number of usable trials per condition was *M =* 33 (*SE* = 1.7), *M =* 38 (*SE* = 1.8), *M =* 35 (SE = 2.6), *M =* 42 (*SE* = 2.3) for Feat, Ortho, Feat-Feat, and Ortho-Ortho, respectively.

#### EEG Recordings

The same EEG system was used to collect data for all of the experiments presented here. A 32-channel sintered Ag/AgC1 electrode array mounted in an elastic cap according to the 10-20 system (FP1, FP2, F3, F4, FZ, F4, F8, FT7, FT8, FC3, FC4, FCz, T3, T4, C3, C4, Cz, TP7, TP8, CP3, CP4, CPz, T5, T6, P3, P4, Pz, O1, O2, and Oz) (QuikCap, Neuroscan Inc.) was used, with the tip of the nose used for a reference. The amplifier was a Neuroscan Synamps with a band pass filter of 0.05-100 Hz and a sampling rate of 500 Hz. The data was reduced offline to 250 Hz. Prior to the start of the experiment, the impedance for all electrodes was lowered to 5 kΩ or less. VEOG and HEOG electrodes were placed on the lower and upper orbital ridge of the left eye as well as the outer canthus of each eye respectively. All ERPs were time locked to the first target onset and a three second epoch (1 second prior to T1 onset, two seconds post) was used for each trial. Baseline activity from a window −200ms to 0ms relative to T1 onset was subtracted from each trial. If the average difference between HEOL and HEOR electrodes in a moving 32ms time window exceeded 20μV, the trial was judged as having contained a horizontal eye movement and was rejected. If the difference between VEOU and VEOL electrodes increases more than 100μV, the trials were marked as having contained a blink and was rejected. Additionally, if any channel exceeded an absolute value of 100μV during the epoched window, that trial was rejected. All artifact rejection and EEG analysis was performed with a combination of custom MATLAB 2012 script and EEGLab 13.3.2b functions (Delorme & Makeig, 2004).

**Figure S5.**
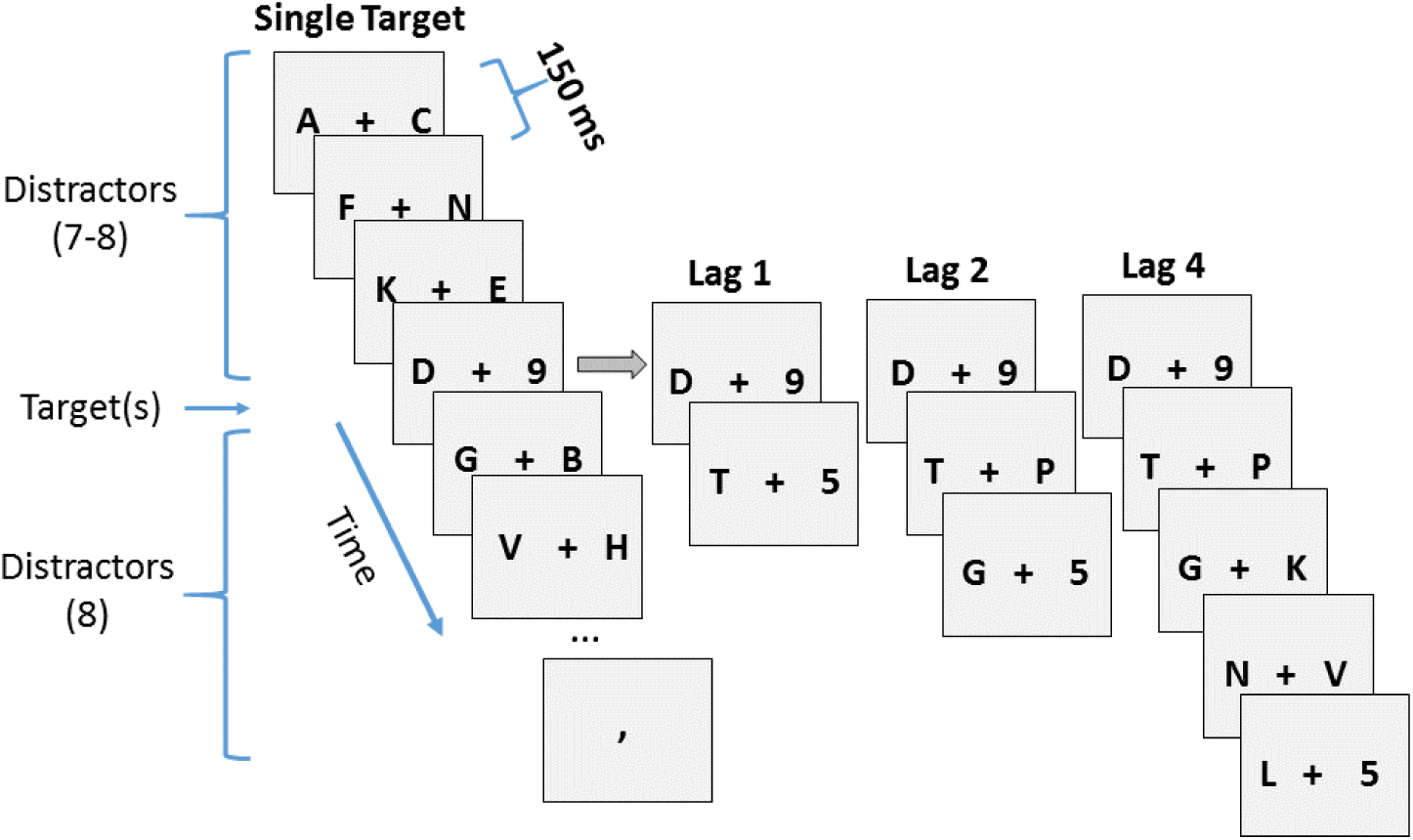
Paradigm for Experiment 3 of Callahan-Flintoft et al. (2018). Participants were presented with 2 RSVP streams. There were 7 or 8 frames of distractors (150ms SOA) prior to target presentation and 8 frames of distractors presented afterwards. Targets were black digits among black letter distractors. T2 was presented either at lag-1, 2, or 4.

## Footnotes

1 We squared AUC values after subtracting them from chance-level because of a unique and interesting implication of below chance-level (or mis-) classification in temporal generalisation plots. Misclassification can imply that the whole-brain pattern difference between classes measured at testing time y resemble those that were present at training time x, *but were of the opposite polarity*, which leads to classifiers making more mistakes than they would by chance.

## REFERENCES

Arlot, S., & Celisse, A. (2010). A survey of cross-validation procedures for model selection. Statistics surveys, 4, 40–79.

Bouthillier, X., Varoquaux, G. (2020) Survey of machine-learning experimental methods at NeurIPS2019 and ICLR2020. [Research Report] Inria Saclay Ile de France. 2020. ffhal-02447823f

Brooks, J. L., Zoumpoulaki, A., & Bowman, H. (2017). Data-driven region-of-interest selection without inflating Type I error rate. Psychophysiology, 54(1), 100–113.

Cawley, G. C., & Talbot, N. L. (2010). On over-fitting in model selection and subsequent selection bias in performance evaluation. The Journal of Machine Learning Research, 11, 2079–2107.

Chikovani, G., Focacci, M. N., Kienzle, W., Lechanoine, C., Levrat, B., Maglic, B., … & Grieder, P. (1967). Evidence for a two-peak structure in the A 2 meson. Physics Letters B, 25(1), 44–47.

Cichy, R. M., Pantazis, D., & Oliva, A. (2014). Resolving human object recognition in space and time. Nature neuroscience, 17(3), 455–462.

Deshpande, G., Li, Z., Santhanam, P., Coles, C. D., Lynch, M. E., Hamann, S., & Hu, X. (2010). Recursive cluster elimination based support vector machine for disease state prediction using resting state functional and effective brain connectivity. PloS one, 5(12), e14277.

Domingos, P. (2012). A few useful things to know about machine learning. Communications of the ACM, 55(10), 78–87.

Dorigo, T. (2015). “Extraordinary claims: the 0.000029% solution”, EPJ Web of Conferences 95, 02003.

Dwork, C., Feldman, V., Hardt, M., Pitassi, T., Reingold, O., & Roth, A. (2015). The reusable holdout: Preserving validity in adaptive data analysis. Science, 349(6248), 636–638.

Eklund, A., Nichols, T., Andersson, M., & Knutsson, H. (2015, April). Empirically investigating the statistical validity of SPM, FSL and AFNI for single subject fMRI analysis. In Biomedical Imaging (ISBI), 2015 IEEE 12th International Symposium on (pp. 1376-1380). IEEE.

Freedman, D. A.(1983). A note on screening regression equations. the American Statistician, 37(2), 152–155.

Geman, S., Bienenstock, E., & Doursat, R. (1992). Neural networks and the bias/variance dilemma. Neural computation, 4(1), 1–58.

Harrison, P.F. (2002). “Blind Analysis”, J. Phys. G: Nucl. Part. Phys. 28, 2679–2691.

Klein, J. R., & Roodman, A. (2005). Blind analysis in nuclear and particle physics. Annu. Rev. Nucl. Part. Sci., 55, 141–163.

King, Jean-Rémi, Alexandre Gramfort, Aaron Schurger, Lionel Naccache, and Stanislas Dehaene. “Two distinct dynamic modes subtend the detection of unexpected sounds.” PloS one 9, no. 1 (2014): e85791.

King, J. R., & Dehaene, S. (2014). Characterizing the dynamics of mental representations: the temporal generalization method. Trends in cognitive sciences, 18(4), 203–210.

Kriegeskorte, N., Simmons, W. K., Bellgowan, P. S., & Baker, C. I. (2009). Circular analysis in systems neuroscience: the dangers of double dipping. Nature neuroscience, 12(5), 535–540.

Kilner, J. M. (2013). Bias in a common EEG and MEG statistical analysis and how to avoid it. Clinical Neurophysiology, 124(10), 2062–3. http://doi.org/10.1016/j.clinph.2013.03.024.

Markoff (2015) Baidu Fires Researcher Tied to Contest Disqualification [Web log post], retrieved from http://bits.blogs.nytimes.com/2015/06/11/baidu-fires-researcher-tied-to-contest-disqualification/

Ng, A. Y. (1997, July). Preventing” overfitting” of cross-validation data. In ICML (Vol. 97, pp. 245–253).

Stone, M. (1974). Cross-validatory choice and assessment of statistical predictions. Journal of the royal statistical society. Series B (Methodological), 111–147.

Varoquaux, G., Raamana, P. R., Engemann, D. A., Hoyos-Idrobo, A., Schwartz, Y., & Thirion, B. (2017). Assessing and tuning brain decoders: cross-validation, caveats, and guidelines. NeuroImage, 145, 166–179.

Vul, E., Harris, C., Winkielman, P., & Pashler, H. (2009). Puzzlingly high correlations in fMRI studies of emotion, personality, and social cognition. Perspectives on psychological science, 4(3), 274–290.

